# Real-Time Visualization of Protein Microenvironment Changes with High Spatial Resolution in Live Cells via Site-Specific Incorporation of Rotor-Based Fluorescent Noncanonical Amino Acids

**DOI:** 10.1101/2024.10.19.619218

**Authors:** Shudan Yang, Shikai Jin, Mengxi Zhang, Yuda Chen, Yiming Guo, Yu Hu, Peter G. Wolynes, Han Xiao

**Affiliations:** Department of Chemistry, Rice University, 6100 Main Street, Houston, TX 77005, USA; Center for Theoretical Biological Physics, Rice University, 6100 Main Street, Houston, TX 77005, USA; Department of Biosciences, Rice University, 6100 Main Street, Houston, TX 77005, USA; Department of Physics, Rice University, 6100 Main Street, Houston, Texas, U.S.A.; Department of Bioengineering, Rice University, 6100 Main Street, Houston, TX 77005, USA; SynthX Center, Rice University, 6100 Main Street, Houston, TX 77005, USA

## Abstract

Traditional methods, such as the use of fluorescent protein fusions and environment-sensitive fluorophores, have limitations when studying protein microenvironment changes at the finest spatial resolution. These techniques often rely on bulky proteins or tags restricted to the N- or C-terminus, which can disrupt the natural behavior of the target protein and dramatically limit the ability of their method to investigate noninvasively microenvironment effects. To overcome these challenges, we have developed an innovative strategy to visualize microenvironment changes of protein substructures in real-time by genetically incorporating environment-sensitive noncanonical amino acids (ncAAs) containing rotor-based fluorophores (RBFs) at specific positions within a protein of interest. Through computational redesign of aminoacyl-tRNA synthetase, we successfully incorporated these rotor-based ncAAs into various proteins in mammalian cells. By site-specifically placing these ncAAs in distinct regions of proteins, we detected microenvironmental changes of several different protein domains during events such as aggregation, clustering, aggregation disassembly, and cluster dissociation.

## Introduction

The protein microenvironments are strongly influenced by physical stress^1–4^, chemical components^5–9^, and many biological factors^10–14^, linked to specific intracellular regulatory pathways^15–18^ and their dysfunction to many human diseases^19–23^. The change of microenvironmental homeostasis of a protein can lead to protein aggregation or clustering, and is often accompanied by abrupt shifts in molecular density, local polarity, and viscosity of protein subdomains, thus allowing detection by environmentally sensitive probes^24–28^. Functional protein cluster typically consists of several functional proteins that respond to the microvariations in the surrounding cytoplasm and their assemble state is closely regulated. In contrast, sometimes, protein aggregation is dysfunctional, involving less malleable intermolecular interactions.^29^ The aggregation and clustering of these proteins can cause significant alterations in the microenvironment of specific protein substructures. Therefore, detecting microenvironmental changes within distinct protein domains, rather than the entire protein, will provide deeper insights into a variety of human diseases and potentially lead to the development of new therapeutic strategies.

Over the past decades, several research tools have been developed to study the overall environment variations of proteins, including proteomic method, based on mass spectrometry ex vivo^30–32^, monitoring in vivo fluorescent proteins combined with biosensors^33–36^, or fluorogenic small molecule probes bound through protein tags. Proteomics-based methods do not allow continuous monitoring of living cells. Using fluorescent proteins to modify targets potentially disrupts the behavior of the target protein in vivo. In addition, the placement of fluorescent proteins or protein tags is often restricted to the N- or C-terminus of target proteins, preventing fluorescent proteins or small probes from detecting the microenvironment changes in spatially distinct regions of the target protein. Thus, a method capable of detecting microenvironmental variations sensitive to different regions of a protein having a high resolution needs to be developed.

Rotor-based fluorophores (RBFs) provide exceptional responses to environmental changes^37–46^. In a low-viscosity solvent, the internal rotation of a π-rich carbon-carbon linkage between electron donor and acceptor of RBFs causes the fluorescence excitation to dissipate non-radiatively. Increasing the viscosity of the environment inhibits the necessary twisted intramolecular charge transfer (TICT), resulting in enhanced fluorescent emission of RBFs in a more rigid environment (**Fig. 1A**).^47,48^ The site-specific installation of an RBF at any desired site of protein will allow for the monitoring of local environmental rigidity in a specific protein domain. To genetically introduce this RBF probe at a desired position of the protein in living cells, Genetic Code Expansion (GCE) Technology is one of the most powerful approaches ^49–53^. GCE technology involves using an aminoacyl-tRNA synthetase/tRNA pair to catalyze the aminoacylation reactions between unnatural amino acid and tRNA, by placing an amber stop codon (TAG) at a specific site in a gene and employing exogenous addition of a noncanonical amino acid (ncAA) to achieve the protein engineering objectives^49,53–55^. Over the past several decades, GCE has been used to produce protein complexes with diverse and valuable properties and functions^56–64^, and thus probably offering powerful tool for system biology.

**Figure 1.**
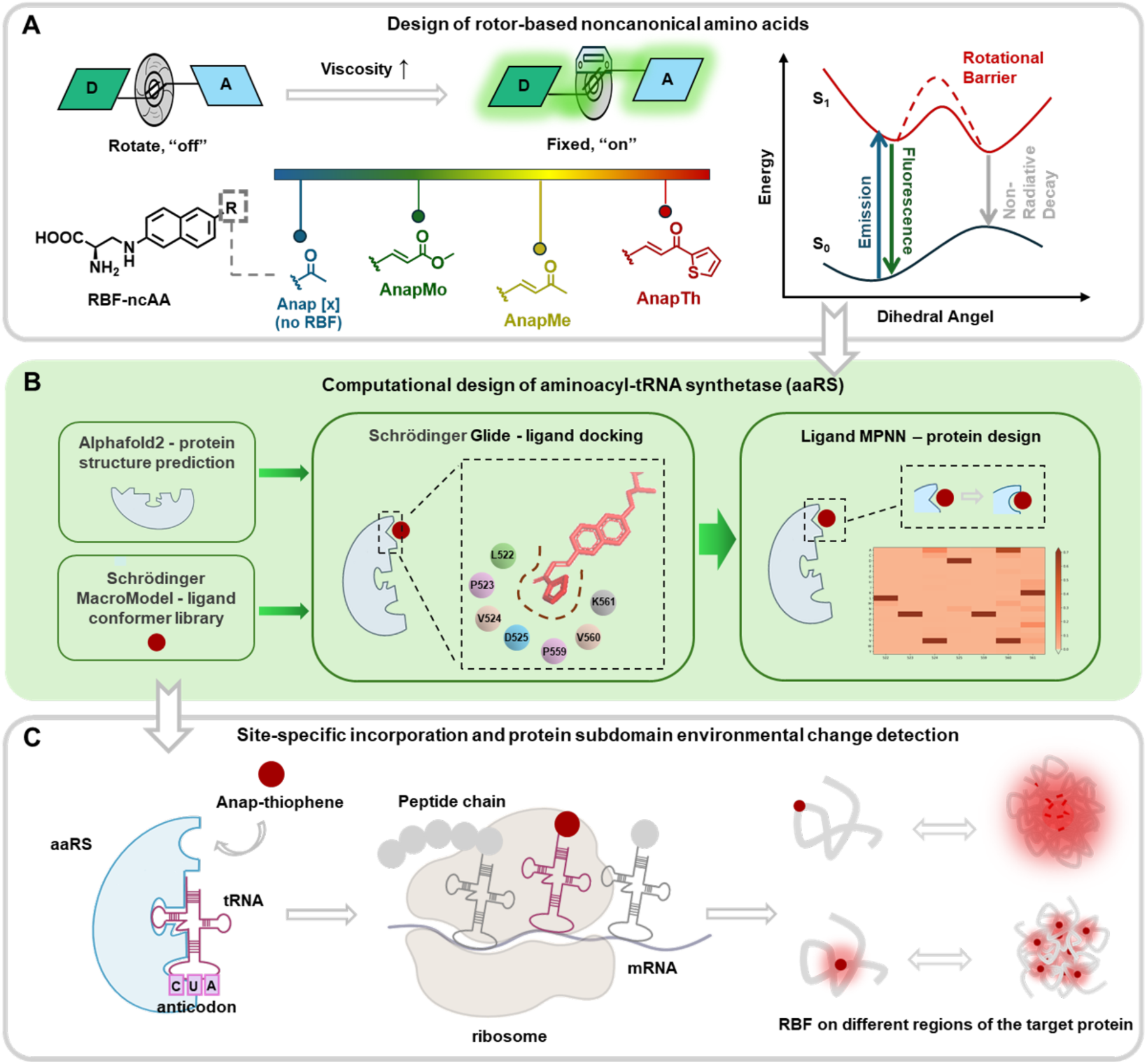
An overview of the development of a novel probe of protein microenvironment changes, combining RBF dyes, computational protein design, and Genetic Code Expansion technology. **(A)** Three viscosity-sensitive noncanonical amino acids were designed based on the Anap skeleton, following the rotor-based mechanism. **(B)** A series of computational methods were employed to redesign Anap-thiophene specific aminoacyl-tRNA synthetase. **(C)** Genetic Code Expansion technology was utilized to site-specifically incorporate Anap-thiophene into the target protein and monitor fluorescence changes with protein conformation.

In this study, we designed a series of rotor-based fluorescent ncAAs by introducing rotational linkage and extending the conjugation system of an Anap ncAA^65^. These synthetic Anap analogs exhibit high sensitivity to the changes of environmental polarity and viscosity (**Fig. 1A**). To apply these novel rotor-based ncAAs for the detection of regional microenvironment changes in proteins, we computationally designed the aaRS to genetically incorporate the variant with the longest emission and largest viscosity factor (AnapTh) into several proteins of interest in a site-specific manner (**Fig. 1B**). With this capability to genetically incorporate AnapTh at a desire position of protein in living cells, we utilized this system to study protein aggregation using Human Superoxide Dismutase (hSOD)and Huntingtin (Htt) as example, and protein clustering, using Stromal Interaction Molecule 1 (STIM1) and Cryptochrome 2 (CRY2). AnapTh located in different domains of each protein exhibits distinct fluorescence behaviors, indicating distinct microenvironmental changes of substructures within the same protein during aggregation or clustering. The technique intellectual in this article provides a general method to explore protein microenvironment changes with a high spatial resolution (**Fig. 1C**).

## Results

### Design of Environmentally Sensitive Noncanonical Amino Acids

Anap, a blue-fluorescent amino acid analog of Prodan^66^, is known for its polarity sensitivity. Due to its small size and favorable quantum yield, Anap is one of the fluorophores that can be genetically incorporated into protein in mammalian cells using GCE technology^65,67,68^. However, its short excitation and emission wavelength (**Fig. 2A** and **2B**), minimal emission shift across different solvent systems (**Fig. 2C**), and barely measurable response to the environmental viscosity changes (**Fig. 2D, 2E** and **2I**) limit its broader applications in real-time fluorescent imaging.

**Figure 2.**
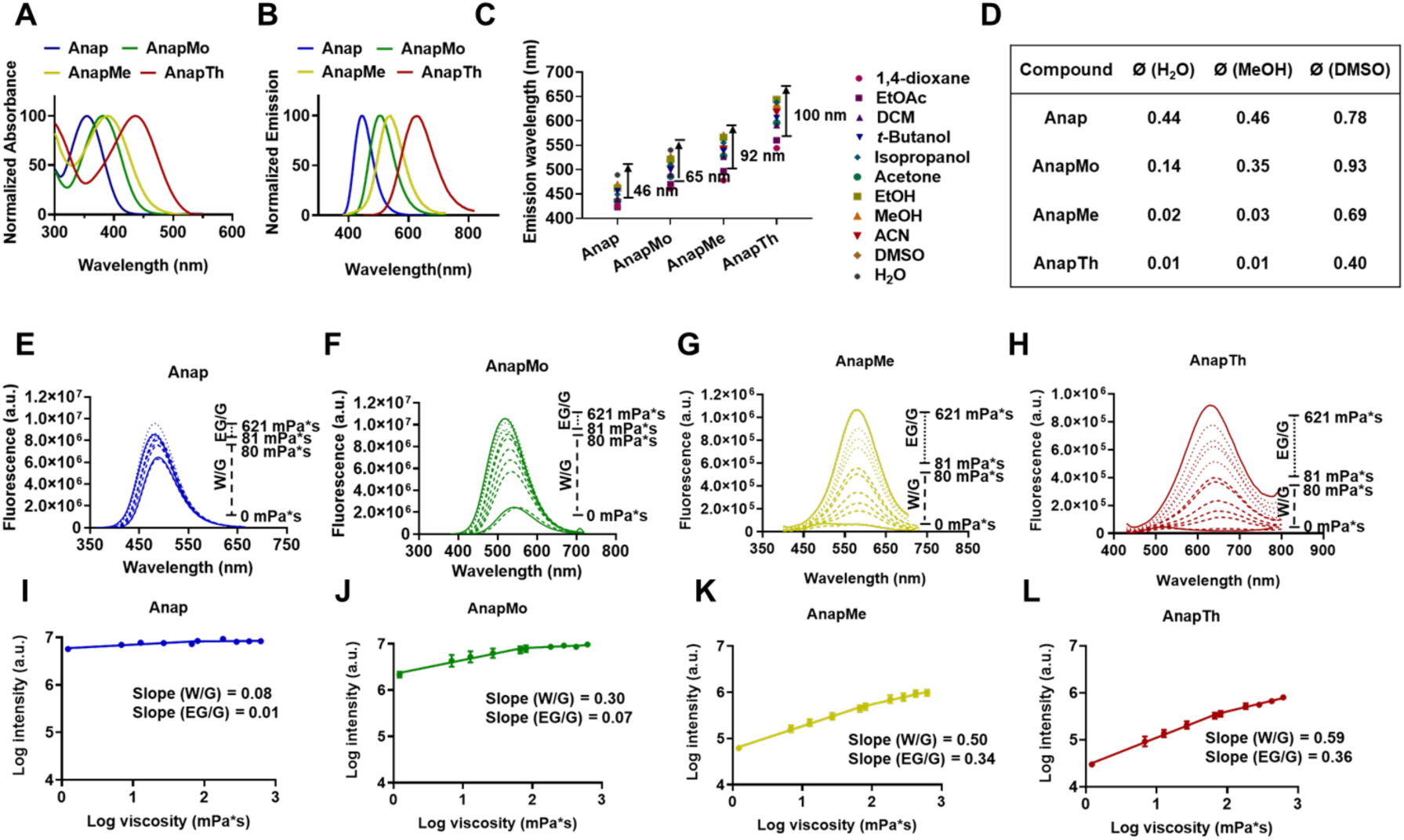
Physical and optical characteristics of four unnatural amino acids: Anap, AnapMe, AnapMo, and AnapTh. **(A)** Absorbance spectrum, **(B)** Emission spectrum, **(C)** Maximum emission wavelength of four noncanonical amino acids in solvents with various polarity, **(D)** Fluorescence quantum yields of four noncanonical amino acids in different solvents. Fluorescence changes of **(E)** Anap, **(F)** AnapMo, **(G)** AnapMe, and **(H)** AnapTh in water/glycerol and glycol/glycerol viscosity gradient systems. Fluorescence intensity – viscosity graphs of **(I)** Anap, **(J)** AnapMo, **(K)** AnapMe, and **(L)** AnapTh in water/glycerol and glycol/glycerol viscosity gradient systems. The slopes of these graphs represent the viscosity sensitivity.

Using Anap as a potential scaffold, we first introduced a rotational linker allowing these rotor-based fluorophores to respond more sensitively to polarity and viscosity. Additionally, we elongated the fluorophore’s emission wavelength by extending the electron conjugation and adjusting its electron-withdrawing groups (EWGs). Inspired by the design of D−A-type naphthalene analogs^69^, we inserted an ethylene unit as a rotor and then conjugated this construct with methoxy-ketone, methyl-ketone, or thiophene-ketone to generate three rotor-based ncAAs called AnapMo, AnapMe and AnapTh (**Fig. 1A**). The ethylene-EWGs were conjugated to the naphthalene through Heck coupling, and the

Misunobu reaction was used to build the amino acid scaffold. All three new ncAAs show more prominent red-shift fluorescence compared to Anap (**Fig. 2A** and **2B**). Their enhanced intramolecular charge transfer (ICT) gives them a larger bathochromic shift of emission wavelength in the solvents with different polarity (**Fig. 2C**). For AnapTh, a difference of the emission shift from the most polar solvent, water, to the non-polar solvent, 1,4-dioxane, is over 100 nm.

As shown in **Fig. 2D**, the ethylene group inserted also affords these ncAAs higher quantum yields in the viscous DMSO when compared to that in non-viscous water and methanol. To further evaluate their viscosity sensitivity, we tested their emission spectra in water/glycerol (0 - 80 mPa*s) and ethylene glycol/glycerol (81 - 621 mPa*s) (**Fig. 2E-H**).

Viscosity sensitivity (x) is a quantitative measure describing the relationship between fluorescence intensity and viscosity^38,70^. Based on the Forster-Hoffmann equation [Eq. (1)],

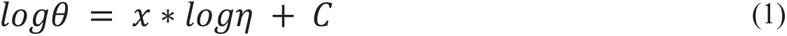

the measured fluorescence intensity maxima (proportional to quantum yield Φ) were plotted against the solvent viscosities in a double logarithmic scale to give a linear correlation, whose slope defines the value of x. Based on our measurements (**Fig. 2I-L**), AnapTh shows the highest viscosity sensitivity in both phases (0.59 in W/G and 0.36 in EG/G). The exceptional sensitivity to both polarity and viscosity, along with the longest emission wavelength of AnapTh, makes it an excellent candidate for studying protein microenvironment variations.

### Computational Design of Aminoacyl-tRNA Synthetase for Genetic Incorporation of AnapTh

Noting the similarity between the scaffolds of Anap and AnapTh, we first performed an expression test using AnapRS/tRNA_CUA_^EcLeu^ created in 2009^65^. After co-transfection of the suppression plasmid pAnap encoding for AnapRS/tRNA_CUA_^EcLeu^ pair and plasmid pEGFP-39TAG encoding Enhanced Green Fluorescent Protein (EGFP) into HEK293T cells, the cells were incubated with AnapTh for 24 hours. The expression level of EGFP-39AnapTh, evaluated by fluorescence intensity using confocal, was pretty low compared to the transfected cells expressing EGFP-39Anap.

To enhance the AnapRS activity for efficiently aminoacylating the tRNA_CUA_^EcLeu^ with AnapTh while excluding endogenous amino acids, we used computational methods to predict the binding pocket and modify the active sites (**Fig. 1B**). First, Alphafold v2.2.0 was used to predict the AnapRS structure, which was mutated from wild-type LeuRS. The top 5 models for the complexes as ranked by their energy values were selected for further protein design. AnapTh-AMP was docked into the predicted AnapRS in Maestro with geometric constraints on its same region of Leu-AMP, which is taken from the Leu-AMP site of *E. coli* leucyl-tRNA synthetase (PDB: 4).

Through analysis of the docking results, we noticed that there were steric conflicts between the ligand and two loops within the protein active pocket, suggesting them as the primary cause of the poor substrate-enzyme recognition and low catalysis efficiency. Seven residues, including Leu522, Pro523, Val524, Asp525, Pro559, Val560, and Lys561 residues within these two loops (**Fig. 3A**), predicted to be within 10Å of the thiophene ring of the ligand, were consequently included in the LigandMPNN computational redesign (**Fig. 3B**). The amino acid with the highest average confidence score for each designed residue (522M, 524A, 525N, 559A, 560A, 561R) was picked out for experimental test (**Fig. 3C**). The pAnap plasmids with single-site mutations were prepared as the first round of evolution. These modified AnapRS mutations were transfected into HEK293T cells together with pEGFP-39TAG. After 24h-incubation following ncAA feeding, the expression level of EGFP was evaluated by EGFP fluorescence intensity using a confocal microscope (**Fig. 3D**). Among the single mutants, pAnap-L522M afforded a 70% higher protein expression level in comparison to the original pAnap plasmid, while we observed only background incorporation in the control group without ncAA feeding. To further improve the efficiency and selectivity, a second round of LigandMPNN calculation was carried out on the basis of pAnap-L522M, and mutated amino acids with the highest confidence score were put into the test as double mutation plasmids. As a result, pAnap-L522M V524A gave a more than three times enhancement of EGFP expression and good selectivity as well (**Fig. 3E** and **3F**). This new plasmid was named pAnap-MA, being used in our protein aggregation study, and expressed modified aaRS was named AnapThRS.

**Figure 3.**
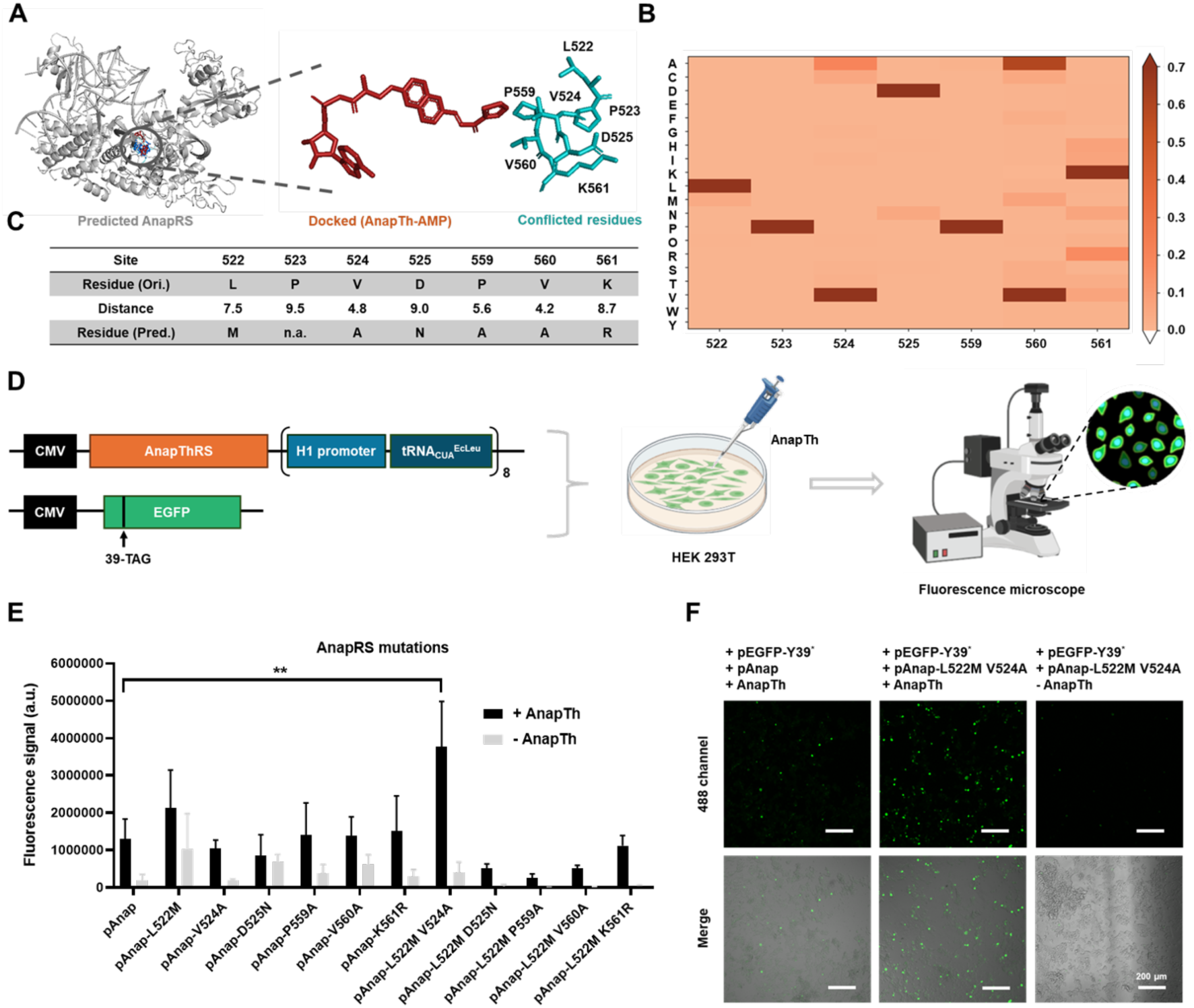
Computaional-assisted aminoacyl-tRNA synthetase. **(A)** Docking of AnapTh-AMP into the AnapRS and the steric conflicts were predicted between the seven residues of AnapRS active pocket and thiophene ring of the ligand. **(B)** The average confidence scores of 20 natural amino acids screened by LigandMPNN. **(C)** The potential mutants of seven conflicted residues. **(D)** The flow chart shows the method to test the working efficiency of AnapRS mutations. AnapRS mutatons and EGFP were cotransfected into HEK293T cells followed by AnapTh feeding. After 24h incubation, the EGFP fluorescence level was estimated by confocal. **(E)** Expression of EGFP-39AnapTh using different AnapRS mutations. The best modification is Leu522M Asp524N. **(F)** Confocal imaging of HEK293T expressing EGFP-39AnapTh with AnapRS and AnapRS 522M 524N mutations. Scale bar = 200 µm. **P≤0.01.

### Study of Human Superoxide Dismutase (hSOD) Microenvironment Changes during Protein Aggregation using AnapTh

Human superoxide dismutase (hSOD) is an essential enzyme for protecting cells from the toxic products of superoxide radicals^71,72^. Its A4V mutation has been reported to be linked to the familial form of amyotrophic lateral sclerosis (FALS), one of the most common neurodegenerative disorders in humans, due to its poor architectural stability and short survival^73,74^. The hSOD-1-A4V mutant protein crystallizes as an asymmetric dimer chelated with Cu/Zn. Protein aggregation initiates from the loss of the metal and leading to the formation of non-metal apo hSOD-1 species, which usually exist as monomers. The metal-free state of hSOD1 tends to assemble into larger aggregated structures under destabilizing conditions^75^ (**Fig. 4A**). During protein aggregation, a reduced local polarity is expected accompanied by an enhanced viscosity because of the exposed hydrophobic protein cores. Since many questions about the role of misfolded and aggregated hSOD-1 in ALS have arisen, we see investigation of hSOD-1 conformational changes should enhance our understanding of the onset and progression of ALS.

**Figure 4.**
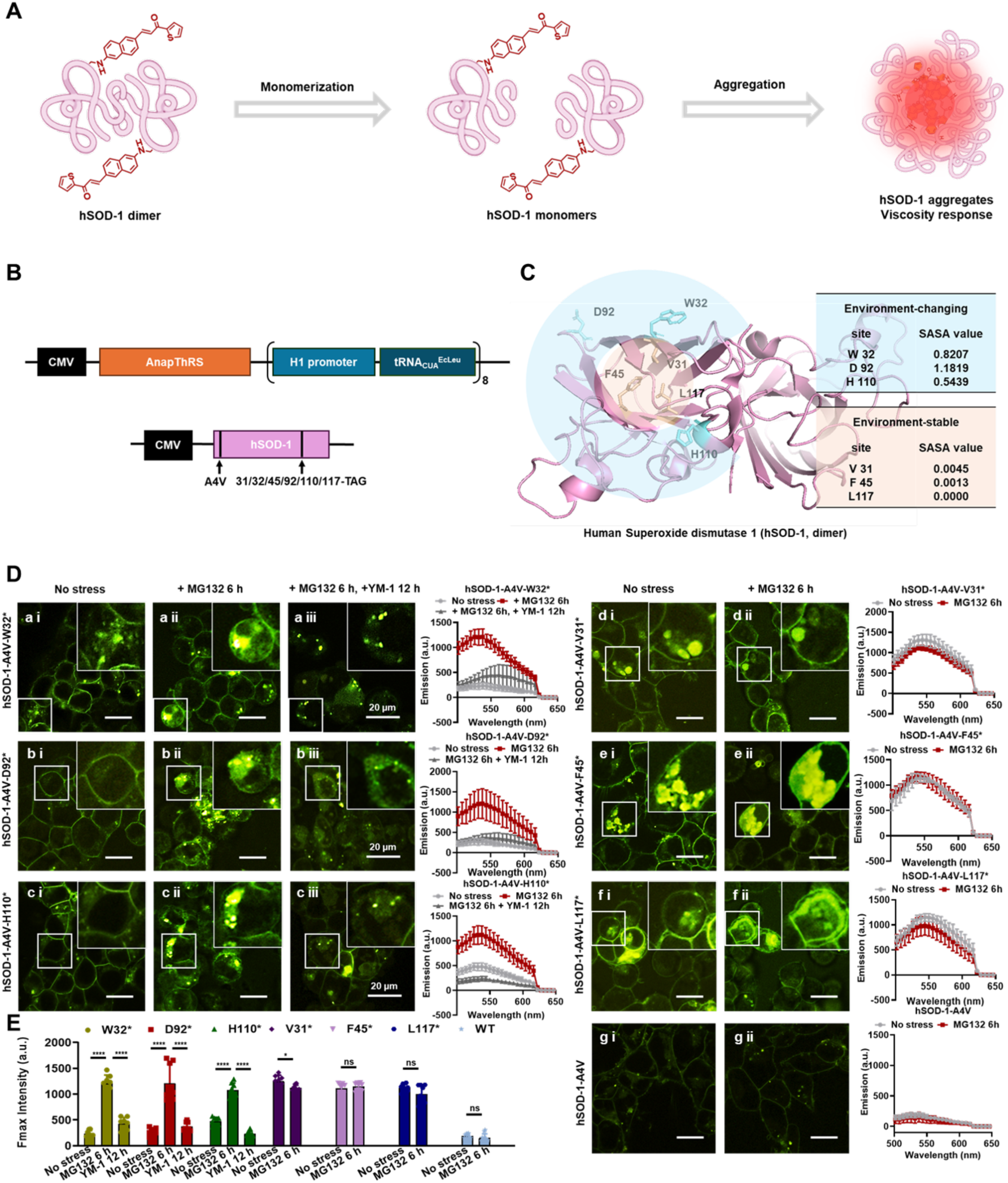
The study of hSOD-1 protein aggregation with AnapTh. **(A)** The scheme of hSOD-1 misfolding and aggregation formation. **(B)** AnapThRS/tRNA_CUA_^EcLeu^ and hSOD-1-A4V plasmids. **(C)** The microenvironment difference between outward residues and interior residues. The TGA ambor code was placed individually at three outward sites with high SASA values: Trp32, Asp92, His110; and three interior sites with low SASA values: Val31, Phe46, Leu117. **(D)** Confocal images of HEK293T cells expressing hSOD-1-A4V with AnapTh inserted at different positions. The fluorescence was scanned by confocal on three stages: no stress, MG132 caused aggregation formation, and YM-1 induced aggregates disassemble. **(E)** Fluorescence intensity summary of AnapTh at different sites on three stages. Scale bar = 20 µm. nsP>0.05, *P≤0.05, **P≤0.01, ****P≤0.0001.

To demonstrate the utility of AnapTh in studying microenvironment changes of hSOD-1-A4V during the aggregation process, we overexpressed hSOD-1-A4V in HEK293T cells with AnapThRS/tRNA_CUA_^EcLeu^ pair. AnapTh was site-specifically incorporated in 6 different positions. Positions Trp32, Asp92, and His110 were located on the protein surface facing outward, in a flexible environment with solvent-accessible surface area (SASA) values close to 1. In contrast, residues at Val31, Phe45, Leu117 were positioned inward in a more stationary environment with the SASA value close to 0 (**Fig. 4B** and **4C**). After 24 hours of post-transfection, cell images were taken, and emission intensities were scanned in the range of 500 nm – 650 nm using Nikon confocal. We found that although a few fluorescent spots in cells expressed hSOD with AnapTh in Trp32, Asp92, and His110 (**Fig. 4D ai, bi** and **ci**) can be observed, they showed fewer counts and exhibited lower brightness than cells with those in Val31, Phe45 and Leu117 (**Fig. 4D di, ei** and **fi**). There was no notable signal from cells transfected with hSOD-1-A4V plasmid without amber stop codon (**Fig. 4D gi**).

We incubated all these groups of cells with MG132 20uM for 6 hours. MG132 is a proteasome inhibitor, causing protein aggregation^76^. After that, more fluorescent puncta with three times higher fluorescent intensity were seen for those cells expressing hSOD-1-A4V with AnapTh incorporated in sites Trp32, Asp92, and His110 (**Fig. 4D aii, bii**, and **cii**). These puncta demonstrate that hSOD-1 coalesces into dense insoluble aggresomes, consistent with previous studies of hSOD aggregation using protein tag-based RBFs. While, in the cells expressing hSOD-1-A4V with AnapTh in sites Val31, Phe45, Leu117, the fluorescence intensity stayed the same and no new punctuated fluorescence appeared (**Fig. 4D dii, eii**, and **fii**). As expected, no fluorescence change was seen in the blank group (**Fig. 4D gii**). Furthermore, by comparing the fluorescence intensities of aii, bii, cii, and di, ei, fi, we see that the viscosity of hSOD interior remains stable during the protein aggregation process (**Fig. 4E**). In the next step, we washed away MG132 and incubated the cells in groups a, b, and c with fresh medium, including 2 µM YM-1. YM-1 is an allosteric Hsp70 inhibitor, which has been reported to reduce aberrant protein aggregation^77^. After 12 hours of YM-1 treatment, we observed that almost punctae fluorescence was diffuse (**Fig. 4D aiii, biii**, and **ciii**) with intensity declination to the nearly initial state (**Fig. 4E**), also combined with a wavelength red-shift that may be due to a polarity change in this process.

### Study of the Fibrillation of Huntingtin (Htt) Using AnapTh

In the hSOD model, we demonstrated that AnapTh performed well in detecting both formation and disassembly of amorphous protein aggregates, as well as distinct fluorescence responses observed across different protein subdomains. To examine the function of AnapTh in different aggregate forms, we shifted to the visualization of a well-studied model of amyloid fiber formation – Huntingtin (Htt) protein. Huntingtin Protein, containing a long repeat of polyglutamine (polyQ) in the N terminus, is well known for its tight association with Huntington’s disease^78,79^. Htt-polyQ fibrils formation occurs through the expansion of CAG triplet repeats in exon 1 of the huntingtin (Htt) gene, forming large polyQ clusters, which also encodes the first 17 amino acids (N-17) that can modulate the aggregation tendency of the expanded polyQ repeat^80,81^. As well as the polyproline at the C terminus, N-17 is subject to many posttranslational modifications that may regulate the function, stability, and distribution of Htt.

In this study, Htt-97Q was overexpressed in HEK293T cells with AnapTh site-specifically incorporated at three different sites, which are located at distances from the poly Q tail, from far to near, as Thr3, Leu7, and Phe11 (**Fig. 5A**). 24 hours after transfection, overexpressed Htt-97Q showed invisible fluorescent dots in the T3 and L7 groups (**Fig. 5B ai** and **bi**), which was observed in the Phe11 group in the cytoplasm (**Fig. 5B ci**). After incubating cells with 20 µM MG132 for 6 hours, for Htt-97Q with AnapTh in Thr3 or Leu7, the diffusive fluorescence observably condensed and formed punctate fluorescent signals within the same cells (**Fig. 5B aii** and **bii**). This observation is consistent with results detected by protein tag-based RBFs, and the fluorescence intensity enhanced three folds (**Fig. 5C** and **5D** from light grey columns to red ones). Unlike Thr3 and Leu7, for the cells expressed Htt-97Q with AnapTh in Phe11, nearly no new fluorescent punctate formed after incubation with MG132, and the fluorescence intensity stayed the same as well (**Fig. 5B cii, 5E**). To explain this observation, we conjectured that the large polyQ moiety affects the environment around residue Phe11, making it more similar to the polarity and viscosity of aggregated protein. After treatment with YM-1, all the punctate fluorescence diffused with intensity declination in Thr3 and Leu7 (**Fig. 5B aiii** and **biii**), and slight wavelength redshift happened in all Thr3, Leu7, and Phe11 cell groups. In addition, we set up the negative control group, cells of which were transfected with Htt-97Q plasmid without TAG, showing extremely low and unchangeable fluorescence during the whole process (**Fig. 5B di, dii, diii**, and **5F**).

**Figure 5.**
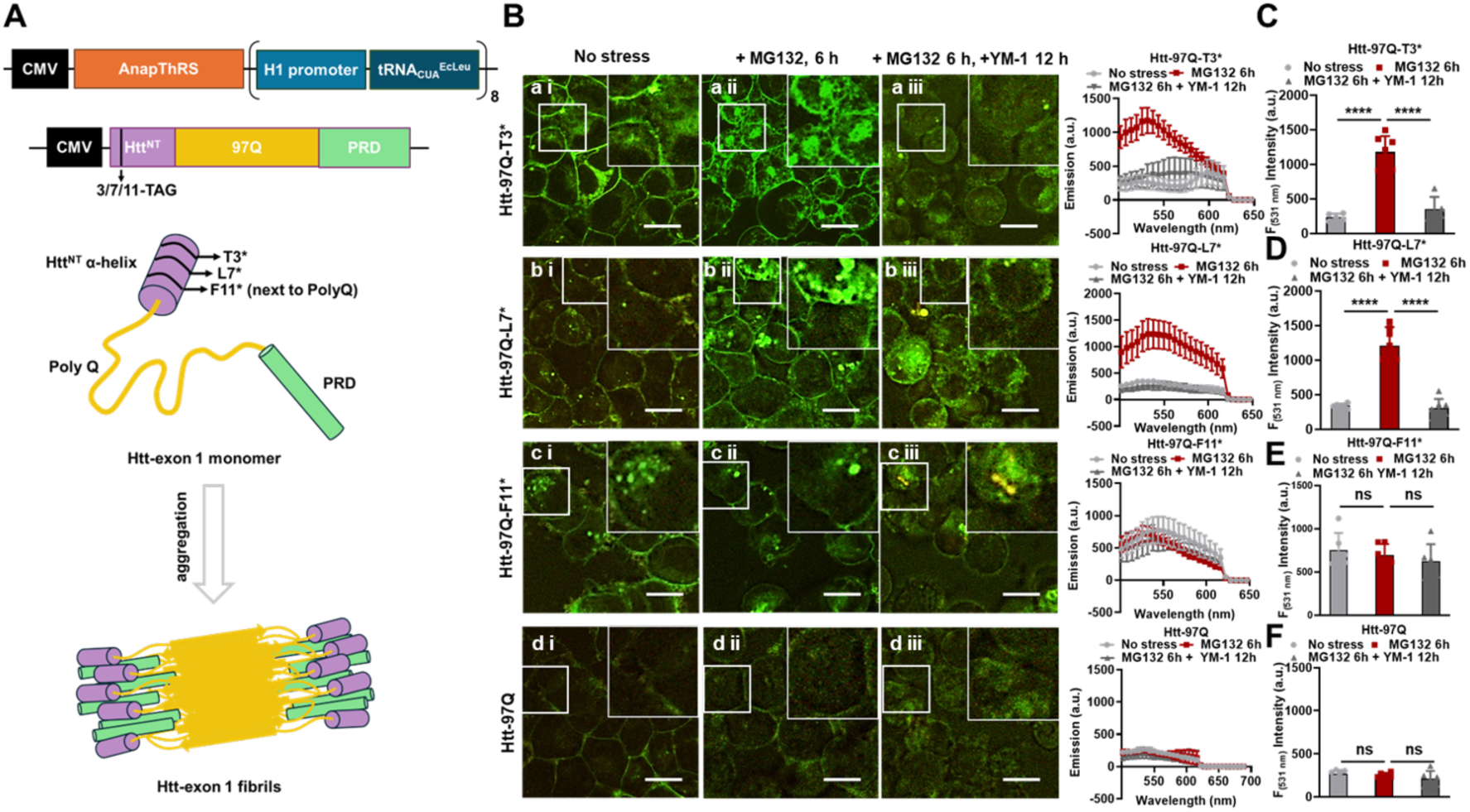
The study of Htt-97Q protein aggregation with AnapTh. **(A)** The scheme of Htt-exon 1 expression through AnapThRS/tRNA_CUA_^EcLeu^ and Htt-97Q plasmids with TGA ambor code placed individually at three different positions: Thr3, Leu7, and Phe11, and fibrils formation in the cytoplasm with drug-induced stress. **(B)** Confocal images of HEK293T cells expressing Htt-97Q with AnapTh inserted at Thr3, Leu7, and Phe11. The fluorescence was scanned by confocal on three stages: no stress, MG132 caused aggregation formation, and YM-1 induced aggregates disassembly. The maximum fluorescence intensity of **(C)** Htt-97Q-3AnapTh, **(D)** Htt-97Q-7AnapTh, **(E)** Htt-97Q-11AnapTh, and **(F)** wild-type Htt-97Q at the starting point, MG132 caused aggregates formation and YM-1 induced aggregates disassembly. nsP>0.05, ****P≤0.0001.

### Tracking of Stromal Interaction Molecule 1 (STIM1) Clustering in Response to Calcium Ion (Ca^2+^) Depletion using AnapTh

To explore if this environmentally sensitive fluorescent ncAA can be applied to study small cluster formation, we introduced Anap-thiophene into a Ca^2+^ sensor protein - STIM1. Calcium (Ca^2+^) is an intracellular secondary messenger that plays important roles in regulating a diverse array of cellular processes. STIM1 is an ER-resident protein, acting pivotally in cellular Ca^2+^ balance regulation. In the inactive state, STIM1 is a dimer with Ca^2+^ binding with ER luminal domains (EF-SAM). When Ca^2+^ is depleted, its dissociation from the EF-hand site triggers the oligomerization of STIM1, inducing puncta formation and delocalization to ER-plasma membrane (PM) junctions in resting cells. The puncta will then activate the PM-resident ORAI Ca^2+^ channels to engage Ca^2+^ influx (**Fig. 6A**).^16,82^

**Figure 6.**
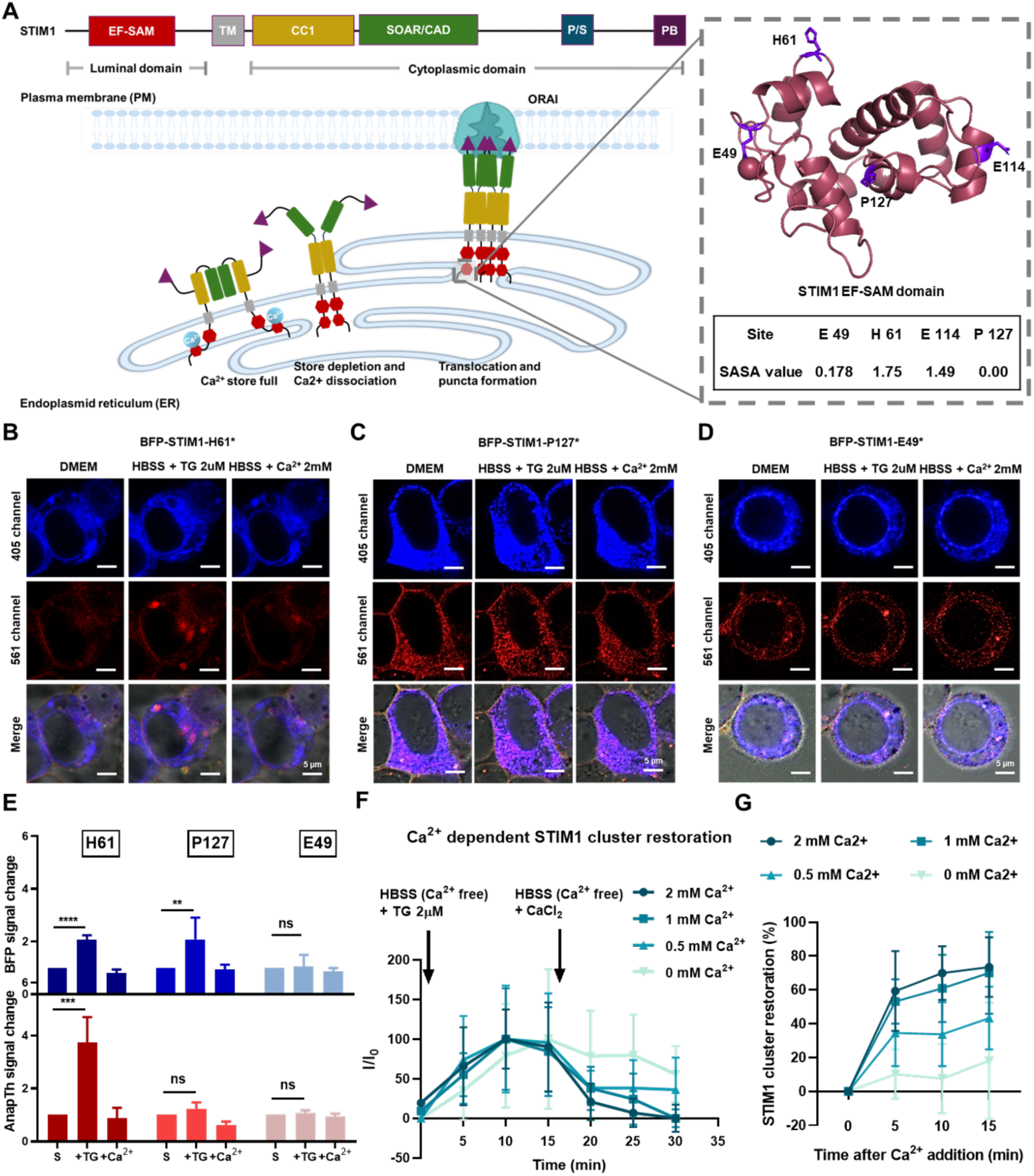
The study of STIM1 protein redistribution and restoration with AnapTh. **(A)** Domain architecture of the human STIM1. SP, signal peptide; EF-SAM, EF-hand, and sterile alpha-motif; TM, transmembrane domain; CC1, coiled-coil domain 1; SOAR, STIM-Orai activating region; P/S, proline/serine-rich region; PB, polybasic tail. The scheme of STIM1– ORAI1 coupling at the ER–PM junction that mediates store-operated Ca^2+^ entry. AnapTh was individually incorporated in Glu49, His61, Glu114, and Pro127 four positions. Confocal images of BFP fused STIM1 with **(B)** His61AnapTh mutation, **(C)** Pro127AnapTh mutation, and **(D)** Glu49AnapTh mutation before and 15 min after 2 µM TG treatment and 15 min after 2 mM CaCl_2_ incubation. **(E)** Comparison of BFP and AnapTh fluorescence changes upon TG or CaCl_2_ with different mutations. **(F)** AnapTh fluorescence of HEK 293T cells expressing BFP-STIM1-His61AnapTh in 15 min of TG treatment and 15 min of restoration in HBSS buffer with Ca^2+^ of different concentrations. **(G)** Kinetic graphs of Ca^2+^ of different concentrations to STIM1 cluster restoration. nsP>0.05, **P≤0.001, ***P≤0.01****P≤0.0001. Scale bar = 5 µm.

To study the microenvironment changes of various regions of EF-SAM during the Ca^2+^ regulation process, we genetically incorporated AnapTh into different sites: His61, which is located on the surface and faces outward; Pro127, which is situated in the domain interior; and Glu49, a residue involved in Ca^2+^ interaction, which served as a control group. In addition, a hypothesis^83^ has been put forward that there must be multiple Ca^2+^ binding sites in EF-SAM. In order to test this hypothesis, we included another residue Glu114, with an intense negative charge (**Fig. 6A**) in the study. Blue fluorescent protein (BFP) was fused with STIM1 to cross-validate the signal changes of AnapTh. The fluorescence of BFP and AnapTh were collected separately through blue and red channels of a Nikon A1 confocal. After treatment with thapsigargin (TG) causing Ca^2+^ store depletion^84^, HEK293T cells expressing BFP fused STIM1 with His61 replaced by AnapTh exhibited fluorescence increasing in both blue and red channels, and bright puncta were apparently observed via the red channel (**Fig. 6B** and **6E**). Despite the redistribution of blue fluorescent protein, the red fluorescence intensity enhancement was much more considerable. After removing the TG and adding 2 mM CaCl_2_ into the HBSS buffer, the fluorescence of both channels returned to the starting level, and the red puncta disappeared. Conversely, the cells expressing BFP-STIM1 with Pro127 replaced by AnapTh exhibited a low blue but high red fluorescence before TG. After incubation with TG, the redistribution and increased signal of BFP indicates the formation of STIM1 clusters, while the red signal remained nearly the same (**Fig. 6C** and **6E**)., supporting that merely elevating the concentration without microenvironment changes does not lead to fluorescence enhancement of RBF dyes. The blue signal was restored after Ca^2+^ was loaded. The Glu49 mutation was expected to result in a loss of one Ca^2+^ binding site on each STIM1 monomer. As shown in **Fig. 6D**, STIM1 oligomers can be observed in the blue channel before TG treatment, while the poor red fluorescence arose from the Glu49 residue not being on the protein surface according to its low SASA (**Fig. 6A**). The subtle effects of TG and Ca^2+^ on BFP distribution and AnapTh and BFP fluorescence intensity indicate the partial loss of the Ca^2+^-sensing region on the EF-SAM domain, and that the Ca^2+^ binding was perturbed (**Fig. 6D** and **6E**). Interestingly, we observed the same phenomenon with cells expressing Glu114 mutation, which supports the hypothesis that there is more than one Ca^2+^-binding region on the EF-SAM domain (**Fig. S6**)^83^.

To further study how the concentration of Ca^2+^ affects the distribution and conformational rearrangement of STIM1, we next measured the Ca^2+^ ion-dependent STIM1 puncta restoration. In this study we employed the HEK293T cells overexpressing BFP-STIM1-H61AnapTh. The starting point of this analysis is to refresh the DMEM to Ca^2+^-free HBSS including 2µM TG. The fluorescence changes of AnapTh were monitored every 5 minutes by Nikon A1 confocal. The highest signal appeared after around 10 minutes (**Fig. 6F**). After 15 minutes, TG-included HBSS was replaced with fresh HBSS with Ca^2+^ of different concentrations (2 mM, 1 mM, 0.5 mM, and 0 mM), and the fluorescence decreasing in different rates was recorded by confocal within 15 minutes. As shown in **Fig. 6F** and **6G**, both 2 mM Ca^2+^ and 1 mM Ca^2+^ stabilized over 80 % of STIM1 within 15 minutes, calculated by the ratio of fluorescence reduction; the rate in 1 mM Ca^2+^ buffer was slightly lower than that in 2 mM Ca^2+^. With lower Ca^2+^ concentrations around 0.5 mM, 40% STIM1 stabilization was completed in 15 minutes. While for cells in an HBSS Ca^2+^-free medium, only a slight fluorescence reduction was detected less than 20%. This Ca^2+^ concentration study provides a new perspective to study the amount of Ca^2+^ binding sites if conducted with all negatively charged residues on EF-SAM domain.

### Tracking of Cryptochrome 2 (CRY2) Optogenetic Clustering Using AnapTh

Cryptochromes (CRY) are photolyase-like blue-light receptors, mediating light responses in both plants and animals. As a critical member of this family, arabidopsis flavoprotein cryptochrome 2 (CRY2) usually interacts with its partner Calcium- and Integrin-Binding protein 1(CIB1) to form reversible complexes.^4,85^ Since these proteins’ interactions do not happen in the dark. As a general optogenetic actuator to manipulate the subcellular processes of protein-protein interactions and a variety of other cellular biochemical events with light, this pair has been attached to diverse target proteins, allowing tight light regulation of target protein activities.^86–88^ On the basis of wild-type CRY2, CRY2clust was first generated by Hyerim Park^89^ through tagging a short peptide, which significantly increases the efficiency of CYR2 clustering. When stimulated by a pulse of blue light, CRY2clust undergoes rapid, reversible, and robust clustering on its own without the help of any partners, redistributing a majority of cytosolic protein into clusters in the cells that are illuminated (**Fig. 7A**). This allows a novel set of experimental approaches for probing cellular biochemistry that existing tools cannot achieve. To incorporate AnapTh into CRY2clust as a small, environmentally sensitive probe for monitoring changes in CRY2 clustering, AnapTh was site-specifically introduced at three different positions (Lys219, Gln263, Pro435) within the full-length CRY2clust using GCE techniques (**Fig. 7B**). HEK293T cells expressing different CRY2clust mutations were stained with NucSpot Live 650 and stimulated with a 488 nm confocal laser with 25% power. Time-dependent red fluorescence changes were recorded through the red channel. Fortunately, the results of all three mutations imply the red fluorescence increases with laser-inducing time and reaches a flat stage in 5 minutes, similar to previous reports. The protein clusters appeared as red dots in the cytoplasm (**Fig. 7C, E**, and **G**). After that, we stopped the laser and kept the cells in the dark for 5 and 10 minutes. The red dots gradually scattered, and the red fluorescence went down to the starting level (**Fig. 7D, F** and **H**).

**Figure 7.**
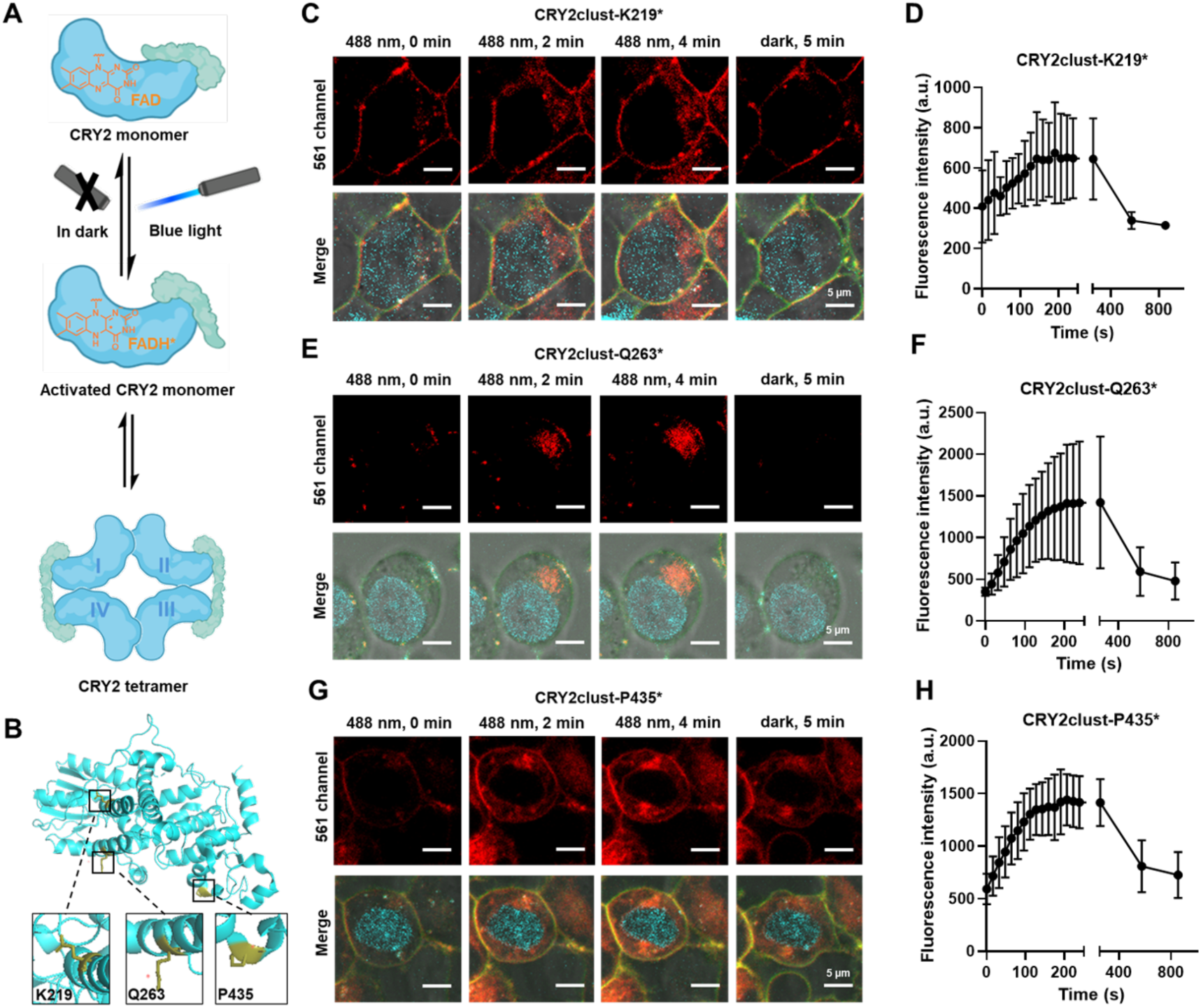
The study of CRY2clust protein clustering with AnapTh. **(A)** Schematic diagram for CRY2clust clustering with light activation. **(B)** AnapTh was individually incorporated into Lys219, Gln263 and Pro435 three positions. Confocal imaging of HEK293T expressing CRY2clust with **(C)** Lys219AnapTh, **(E)** Gln263AnapTh and **(G)** Pro435AnapTh mutation activated by confocal 488nm laser for 2 and 4 minutes and kept dark for another 5 minutes after activation. Fluorescence intensity changes of HEK 293T expressing CRY2clust with **(D)** Lys219AnapTh, **(F)** Gln263AnapTh and **(H)** Pro435AnapTh mutation upon stimulating by confocal 488nm laser for 5 minutes and keeping in dark for another 5 and 10 minutes. Scale bar = 5 µm.

## Discussion

In this work, we developed a new strategy to visualize the microenvironment variations of protein subdomains with spatial resolution in live cells. Although rotor-based fluorogenic probes have been used as proteome stress sensors in previous studies, previous to this work, there was no technology to site-specifically introduce them at desired positions within proteins. This limitation in probe placement prevents spatial-resolution studies of different protein regions and introduces potential errors in the fluorescence responses of the probes to environmental changes within protein subdomains. To overcome these position limitation, we designed the first rotor-based fluorescent noncanonical amino acids AnapTh and showed its site-specific intallation at desired positions of proteins in live cells with the GCE technology. This genetically incorporated AnapTh can replace the fluorescent proteins or protein tags used in previous protein labeling methods, minimizing the impact of the labelling probes on the natural behaviors and functions of target proteins. Moreover, we can now place RBFs in any desired position on target proteins, enabling us to study the microenvironment changes of different regions of the protein during aggregation or clustering processes.

To demonstrate the utility of the approach, we investigated the microenvironmental changes of four protein systems, containing hSOD, Htt, STIM1, and CRY2, which either reversibly form aggregates or form clusters of varying shapes and sizes in response to different stimuli. The results confirm the broad applicability of our rotor-based AnapTh probe. As we know, failure to control biomolecule microenvironments may lead to serious disease. We anticipate that the present fluorogenic proteome sensor system will make possible a wide range of proteostasis studies, illuminate the relationships between the protein microenvironment changes and function at a high spatial resolution, and give us deeper insights into large scale biomolecular assemblies.

## Methods

### General information

Chemical reagents were purchased from Sigma-Aldrich, Oakwood Chemical, TCI Chemicals, and used without further purification. LB agar and 2YT were ordered from BD Difco TM . Isopropyl-β-D-thiogalactoside (IPTG) was purchased from Anatrace. 4-12% Bis-Tris gels for SDS-PAGE were purchased from Invitrogen. QuickChange Lightning Multi Site-Directed Mutagenesis Kit was purchased from Agilent Technologies (Cat. 210515). Oligonucleotide primers were purchased from Integrated DNA Technologies and Eurofins Genomics. Plasmid DNA preparation was carried out with the GenCatch TM Plus Plasmid DNA. Miniprep Kit and GenCatch TM Advanced Gel Extraction Kit. FastBreak™ Cell Lysis Reagent 10x was purchased from Promega (Cat. V8572). Ni-NTA Agarose was obtained from Qiagen (Cat. 30230). 1H and 13C NMR spectra were measured on a Bruker AVANCE III HD 600 MHz spectrometer. UV-vis absorption spectra were measured on a Thermo Scientific Evolution 220 UV-Vis Spectrophotometer. Fluorescence measurements were conducted on a Horiba FluoroMax-4 spectrofluorometer. ESI mass spectroscopy was performed on a Bruker MicroToF ESI LC-MS System. Confocal fluorescent images were performed using Nikon A1R-si Laser Scanning Confocal Microscope (Japan), equipped with lasers of 405/488/561/638 nm.

### Absorbance spectra measurement

Absorbance spectra were collected using a Thermo Scientific Evolution 220 UV-Vis Spectrophotometer. Fluorophores were prepared with a final concentration of 5 µM in DMSO at room temperature and transferred in a 1 cm quartz cuvette for measurements.

### Fluorescence spectra measurement

Fluorescence spectra were recorded using a Horiba FluoroMax-4 spectrofluorometer. Fluorophores were prepared at room temperature with a final concentration of 5 µM in DMSO and transferred to a 1 cm quartz cuvette for measurements.

### Quantum yield determination

Quantum yield measurements were determined using fluorescein (fluorescence quantum yield of 0.92 in 0.1 M NaOH) and Quinine sulfate (fluorescence quantum yield of 0.546 in H2SO4 0.5 M). The fluorescence quantum yield, Φf (sample), was calculated according to the equation as follows:

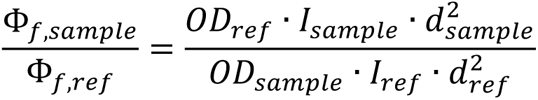

Φ_f_: quantum yield of fluorescence.

I: integrated emission intensity.

OD: optical density at the excitation wavelength.

d: refractive index of solvents, d_CH3CN_=1.34; d_ethanol_=1.36; d_water_=1.33.

### Polarity sensitivity measurement

Four unnatural amino acids are diluted from 50 mM DMSO stock into a final concentration of 25 µM with different solvents at room temperature. The compounds were excited at their λ_abs_, and the fluorescence was collected via Horiba FluoroMax-4 spectrofluorometer. The polarity of each solvent is shown in Table S1.

### Viscosity sensitivity measurement

Four unnatural amino acids are diluted from 50 mM DMSO stock into a series of water/glycerol mixture and ethylene glycol/glycerol mixture at 20 °C with a final concentration of 25 µM. The composition and viscosity of each mixture at 20 °C are described in Table S2.

### Computational protein design

AnapRS was evoluted from LeuRS by Hyun Soo Lee in 2009 with 10 mutations from the reference sequence P07813 in Uniprot: Leu38F, Met40G, Leu41P, Tyr499V, Tyr500L, Tyr527A, His537E, Leu538S, Phe541C, Ala560V. The RMSD between the two structures is 3.07. Alphafold v2.2.0 was used to predict the AnapRS structure. The SMILES format of AnapTh-AMP was drawn in ChemDraw to generate the mol2 format of the 3D structure and aligned to Leu-AMP with the identical portion in 4. After the initial AnapTh-AMP structure was generated, the MacroModel conformational search tool implemented in Schrödinger Suite Release 2018-4 was used to construct the ligand conformational library with strong dihedral constraints force constant 100, degree range 10, torsion value is the same as the Leu-AMP to the bonds that exist in Leu-AMP. Then, the Glide Receptor Grid Generation tool implemented in Schrödinger Suite Release 2018-4 was utilized to generate the receptor grid, which can assist in the identification of the anticipated open volume that a ligand could fit in. The reference ligand position is selected as the original Leu-AMP position. All parameters are set as the default. After that, the Glide Docking tool was used to do the ligand docking. The force field used for docking is OPLS3 and all parameters are set as the default. The receptor grid and the ligand conformational library were imported. The top 5 complex structures that ranked by the energy values were selected for the protein design.

### Plasmid construction

All the plasmids were constructed by PCR and Gibson assembly. Oligonucleotides used in this study are listed in Table S3. Gene fragments used in this study were ordered as g-blocks from Twist Bioscience. DNA sequences are listed in Table S4. Gibson products were transformed into DH10B or Stbl3 (only for pHTN-Htt-97Q construction) strains for amplification.

- For plasmid pcDNA3.1-AnapRS, g-block of AnapRS and tRNACUAEcLeu cassette (construct refer to previous work from Schultz lab1) were cloned into pcDNA3.1 vector (with primer da818, da821).
- Plasmid pcDNA3.1-Anap-Mutants were generated from pcDNA3.1-AnapRS by single-site mutation. Primer used as listed in Table S3.
- Plasmid pEGFP39TAG was a gift from Jiantao Guo’s lab2.
- For plasmid pHTN-Htt-97Q, Htt-97Q were subcloned from addgene plasmid #1184 (Htt-97Q) with primer mz493, mz494. pHTN vector was subcloned from plasmid pHTN-Halo constructed by our lab in previous work3 with primer mz495, mz497. PCR products then were connected via Gibson assembly. All the plasmid with TAG mutation generated from pHTN-Htt-97Q by single-site mutation. Primer used as listed in Supplementary Table S3.
- For plasmid pHTN-hSOD1-A4V, g-block of hSDO1-A4V was amplified with primer mz498, mz499. pHTN vector was subcloned from plasmid pHTN-Halo constructed by our lab in previous work3 with primer mz500, mz502. PCR products then were connected via Gibson assembly. All the plasmids with TAG mutation were generated from pHTN-hSOD1-A4V by single-site mutation. Primer used as listed in Supplementary Table S3.
- For plasmid pCMV6-XL5-tagBFP-STIM1, tagBFP was subcloned from addgene plasmid #75160 with primer mz707, mz708. pCMV6-XL5-STIM1 was subcloned from a gift plasmid pCMV6-XL5-EGFP-STIM1 given by Yubin Zhou Lab^83^ with primer mz709, mz710. PCR products then were connected via Gibson assembly. All the plasmids with TAG mutation were generated from pCMV6-XL5-tagBFP-STIM1 by single-site mutation. Primer used as listed in Supplementary Table S3.
- For plasmid pcDNA3.1-BFP-CRY2-3xSGG-tRNA, tagBFP was subcloned from addgene plasmid #75160 with primer mz711, mz712. pcDNA3.1-tRNA was subcloned from pcDNA3.1-AnapRS mentioned above with primer mz641, mz642. CRY2-3xSGG was subcloned from a gift plasmid mCh-CRY2clust-Capture 2.0 given by Yubin Zhou Lab^86^ with primer mz662, mz663. PCR products then were connected via Gibson assembly. All the plasmids with TAG mutation were generated from pcDNA3.1-BFP-CRY2-3xSGG-tRNA by single-site mutation. Primer used as listed in Supplementary Table S3.

### Transfection and live cell imaging

HEK293T cells were seeded at ∼25% confluency in Lab-Tek 8-chamber #1.0 Borosilicate coverglass system or 35 mm glass bottom dish (poly-D-lysine treated, MetTek Corporation) in Dulbecco’s modified Eagle’s medium (DMEM, Gibco) supplemented with 10% Fetal Bovine Serum (FBS, Gibco) and 0.5 % mg/mL penicillin-streptomycin-glutamine (PSQ, Gibco). When cells reached 60% confluency, transfection was carried out using PolyJetTM in vitro DNA transfection reagent. After that, HEK293T cells were fed with Anap-thiophene and incubated for 48 h. After 48 h, excess ncAA was washed away by replacing the DMEM media three times prior to confocal microscopy experiments. Live cell imaging was carried out using a Nikon Instruments A1 Confocal Laser Microscope with a 60xoil lens at room temperature. Results were analyzed using software NIS-Element Analysis and NIS-Element Viewer.

### Small molecule-induced aggregation and disaggregation

MG132: For HEK 293T cells expressing wild-type or mutated hSOD-1-A4V or Htt-97Q, excess ncAA was washed away by fresh DMEM full medium two times, and then cells were incubated in fresh DMEM with MG132 20 µM to induce proteome aggregation. Cells were treated for 6 to 8 hours prior to fluorescence confocal microscopy.

YM-1: DMEM medium with MG132 was replaced by fresh medium with 2 µM YM-1. Cells were incubated for 12 hours, and subsequently excess YM-1 was washed away by fresh DMEM medium two times prior to the fluorescence confocal microscopy.

### Ca^2+^ disassociation induced STIM1 redistribution

For HEK293T cells expressing BFP-STIM1 mutations, DMEM medium was replaced by prewarmed Ca^2+^-free Hank’s balanced salt solution (HBSS), including 2 µM thapsigargin used to trigger store depletion. Live cell imaging was performed after 5, 10, and 15 minutes. After that, thapsigargin was removed by replacing the medium with HBSS with CaCl2 of gradient concentrations. Live cell imaging was performed after 5, 10, and 15 minutes.

### Light-induced CRY2clust clustering

For HEK293T cells expressing CRY2clust mutations, the excess nuclear dye was washed away by fresh DMEM two times. Photoactivation was delivered with a 488 nm Nikon A1 confocal laser with 25% maximum laser power. Live cell imaging was collected during the whole illumination process. After 5 minutes of illumination, laser scanning was stopped, and images were taken at 5 and 10 minutes.

### Nuclear stain

For HEK293T cells expressing CRY2clust mutations, DMEM medium was replaced by fresh DMEM with diluted 1000x NucSpot Live 650 DMSO stock solution. After incubation for 20 to 30 minutes, excess nuclear dye was washed away by fresh DMEM two times.

### Organic synthesis of four noncanonical amino acids

All schemes, procedures, and identifications are available in Supporting information.

## Acknowledgments

We thank Dr. Xiao Laboratory members for insightful comments. This work was supported by SynthX Seed Award (SYN-IN-2024-002), NIH (R35-GM133706, R01-CA277838,, and R01-AI165079 to H.X.), the Robert A. Welch Foundation (C-1970 to H.X.), US Department of Defense (W81XWH-21-1-0789 and T9425-23-1-0494 to H.X.), Center for Theroretical Biological Physics (NSF grant PHY-2019745 to P.G.W.) and D. R. Bullard Welch Chair at Rice University (Grant C-0016 to P.G.W). H.X. is a Cancer Prevention & Research Institute of Texas (CPRIT) scholar in cancer research.

