## Supporting Information for "Real-Time Visualization of Protein Microenvironment Changes with High Spatial Resolution in Live Cells via Site-Specific Incorporation of Rotor-Based Fluorescent Noncanonical Amino Acids"

### Synthesis of non-canonical amino acids

#### L-3-(6-acetylnaphthalen-2-ylamino)-2-aminopropanoic acid (Anap)

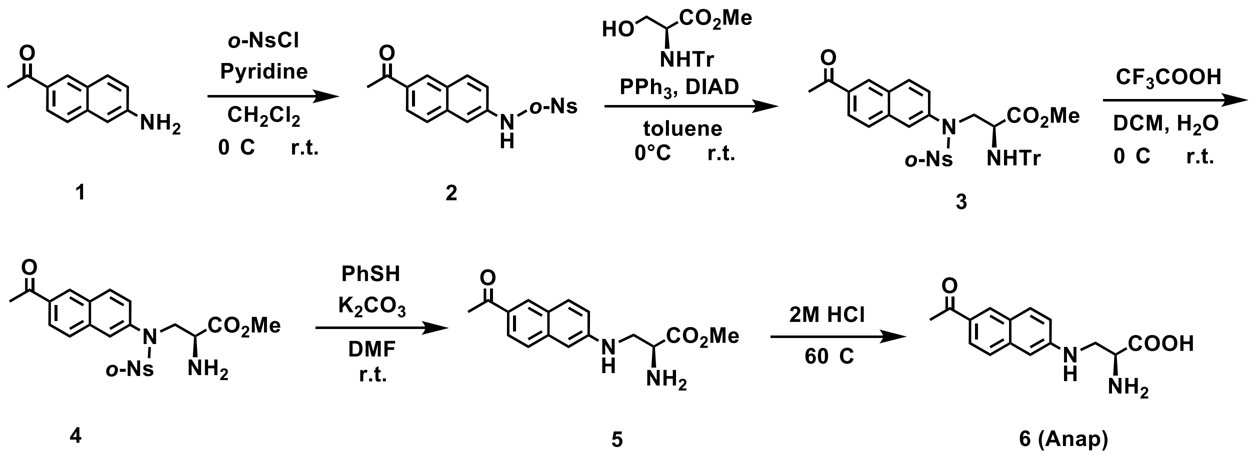

The synthesis of **Anap** followed the method reported in Zheng Xiang and Lei Wang[1].

To a solution of compound **1** (50 mg, 1 eq) in $CH_2Cl_2$ (3mL) were added pyridine (0.024 mL, 1.1 eq) and *o*-NsCl (62 mg, 1.05 eq) sequentially at 0 °C. The reaction was stirred at room temperature overnight. The reaction mixture was washed with 1 M HCl solution, water, and brine and dried over $Na_2SO_4$. The solution was concentrated, and the resulting red solid **2** was dissolved in toluene (3 mL) and stirred at 0 °C. To this mixture were added N-trityl-L-serine methyl ester (195 mg, 2 eq) and $PPh_3$ (142 mg, 2 eq). DIAD (0.11 mL, 2 eq) was added dropwise to the solution. The reaction mixture was stirred overnight at room temperature. The solution was concentrated and purified by silica chromatography to give compound **3** (150 mg, 78%) as a red solid.

To a solution of **3** (100 mg, 1 eq) in $CH_2Cl_2$ (5 mL) were added TFA (1 mL) and water (1 mL) at 0 °C. The reaction mixture was stirred at room temperature for 3 h and concentrated under vacuum. The residue was dissolved in DMF (1 mL). Thiophenol (31 mg, 2 eq) and $K_2CO_3$ (194 mg, 10 eq) were added to the solution sequentially. The reaction mixture was stirred at room temperature for 2 h. Water was added to the mixture, and the solution was extracted with EtOAc three times. The combined organic phase was washed with brine, dried over $Na_2SO_4$, and filtered. The filtrate was concentrated and purified by silica chromatography to give compound **5** (30 mg, 72%) as yellow solid.

A solution of compound **5** (30 mg, 1 eq) in 2 M HCl solution (1 mL) was stirred at 60 °C for 5 h. The reaction mixture was lyophilized to give a yellow solid. After being washed with diethyl ether, the solid was dried under vacuum to give **Anap** (28 mg, 99%) as a yellow solid. $^1$H NMR (600 MHz, D2O) δ 8.32 (s, 1H), 7.81 – 7.77 (m, 2H), 7.67 (d, J = 8.7 Hz, 1H), 7.64 – 7.58 (m, 1H), 7.12 (dd, J = 8.9, 2.2 Hz, 1H), 7.01 (d, J = 1.9 Hz, 1H), 3.65 – 3.60 (m, 2H), 3.39 (td, J = 10.1, 4.4 Hz, 1H), 2.67 (s, 3H). $^{13}$C NMR (151 MHz, D2O) δ 203.26, 180.99, 148.90, 138.09, 131.62, 131.18,

129.72, 125.95, 125.56, 124.15, 119.08, 103.78, 55.01, 47.43, 25.77. HRMS $[M+H]^+$ calcd. for $C_{15}H_{17}N_2O_3$ 273.1239, found 273.1241.

#### Methyl-L-3-(6-((2-amino-3-methoxy-3-oxopropyl)amino)naphthalen-2-yl)acrylate (AnapMo)

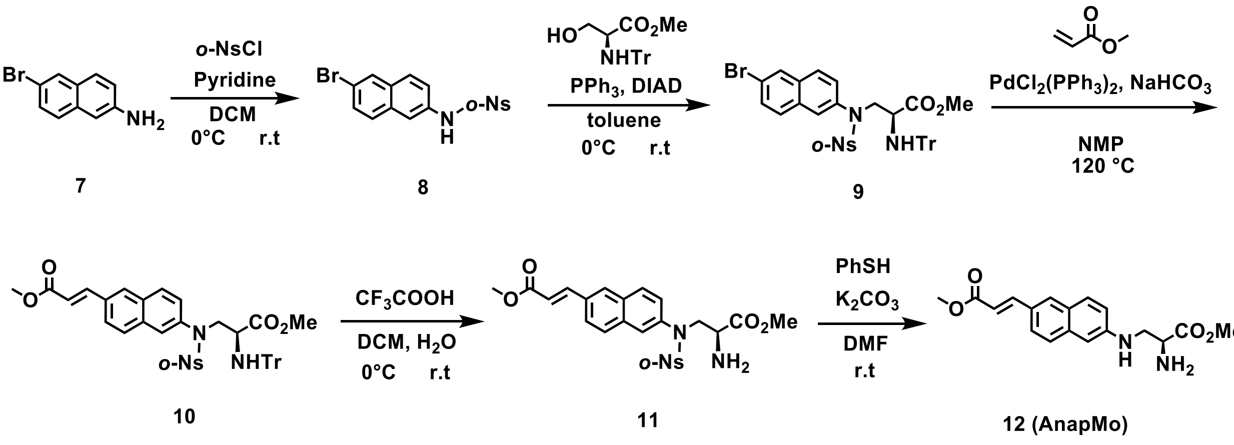

To a solution of compound **7** (1 g, 1 eq) in $CH_2Cl_2$ (45 mL) were added pyridine (0.4 mL, 1.1 eq) and *o*-NsCl (1.05 g, 1.05 eq) sequentially at 0 °C. The reaction was stirred at room temperature overnight. The reaction mixture was washed with 1 M HCl solution, water, and brine and dried over $Na_2SO_4$. The solution was concentrated, and the resulting crude product was purified through silica chromatography to give **8** (1.2 g, 63%). $^1H$ NMR (600 MHz, $CDCl_3$) δ 7.94 – 7.93 (m, 1H), 7.86 (dd, J = 8.0, 1.1 Hz, 1H), 7.82 (dd, J = 7.9, 1.3 Hz, 1H), 7.68 – 7.61 (m, 4H), 7.55 – 7.51 (m, 2H), 7.36 (dd, J = 8.8, 2.2 Hz, 1H).

To a solution of compound **8** (500 mg, 1 eq) in toluene (10 mL) and stirred at 0 °C. To this mixture were added N-trityl-L-serine methyl ester (842 mg, 2 eq) and $PPh_3$ (611 mg, 2 eq). DIAD (0.46 mL, 2 eq) was added dropwise to the solution. The reaction mixture was stirred overnight at room temperature. The solution was concentrated and purified by silica chromatography to give compound **9** (655 mg, 75%). $^1H$ NMR (600 MHz, $CDCl_3$) δ 7.99 (d, J = 1.1 Hz, 1H), 7.82 (d, J = 1.6 Hz, 1H), 7.67 (dd, J = 8.8, 3.1 Hz, 2H), 7.63 – 7.57 (m, 3H), 7.38 (ddd, J = 10.9, 8.4, 1.5 Hz, 2H), 7.35 – 7.31 (m, 1H), 7.27 (dd, J = 7.1, 2.3 Hz, 6H), 7.13 – 7.07 (m, 9H), 4.33 (dd, J = 14.5, 4.2 Hz, 1H), 4.30 – 4.24 (m, 1H), 3.56 – 3.48 (m, 1H), 3.17 (s, 3H), 2.64 (d, J = 10.3 Hz, 1H).

Compound **9** (100 mg, 1eq), $PdCl_2(PPh_3)_2$ (4.6 mg, 0.05 eq), and $NaHCO_3$ (22 mg, 2 eq) were contained in a reaction flask with nitrogen protection, and anhydrous NMP (2.5 mL) was added. Then, methyl acrylate (0.021 mL, 1.7 eq) was added. The reaction flask was sealed and stirred at 120 °C overnight. The mixture was then cooled to room temperature, diluted with EA, and washed with brine. The organic layer was dried over anhydrous $Na_2SO_4$, filtered, and condensed under reduced pressure. The crude product was purified by silica-gel flash column chromatography to obtain the desired product **10** (51 mg, 52%). $^1H$ NMR (600 MHz, CDCl3) δ 7.90 (s, 1H), 7.85 (dd, J = 8.8, 7.1 Hz, 2H), 7.78 (t, J = 8.2 Hz, 2H), 7.70 (dd, J = 8.6, 1.3 Hz, 1H), 7.63 – 7.56 (m, 2H), 7.42 (dd, J = 8.0, 1.0 Hz, 1H), 7.38 (dd, J = 8.8, 2.0 Hz, 1H), 7.35 – 7.31 (m, 1H), 7.25 (d, J = 1.3 Hz, 4H), 7.12 – 7.05 (m, 9H), 6.58 (d, J = 16.0 Hz, 2H), 4.34 (dd, J = 14.6,

4.1 Hz, 1H), 4.28 (dd, J = 14.5, 6.9 Hz, 1H), 3.83 (d, J = 17.5 Hz, 3H), 3.57 – 3.50 (m, 1H), 3.17 (d, J = 13.4 Hz, 3H), 2.63 (d, J = 10.3 Hz, 1H).

To a solution of **10** (50 mg, 1 eq) in $CH_2Cl_2$ (5 mL) were added TFA (1 mL) and water (1 mL) at 0 °C. The reaction mixture was stirred at room temperature for 3 h and concentrated under vacuum. The residue was dissolved in DMF (1 mL). Thiophenol (16 mg, 2 eq) and $K_2CO_3$ (99 mg, 10 eq) were added to the solution sequentially. The reaction mixture was stirred at room temperature for 2 h. Water was added to the mixture, and the solution was extracted with EtOAc three times. The combined organic phase was washed with brine, dried over $Na_2SO_4$, and filtered. The filtrate was concentrated and purified by silica chromatography to give compound **12 AnapMO** (20 mg, 85%) as yellow solid. $^1H$ NMR (600 MHz, $CDCl_3$_3mm_tubes) δ 7.79 (d, J = 15.9 Hz, 1H), 7.74 (s, 1H), 7.62 (d, J = 8.8 Hz, 1H), 7.56 (q, J = 8.6 Hz, 2H), 6.92 (dd, J = 8.8, 1.6 Hz, 1H), 6.83 (s, 1H), 6.44 (d, J = 15.9 Hz, 1H), 4.72 – 4.58 (m, 1H), 3.82 (d, J = 16.8 Hz, 3H), 3.77 (s, 3H), 3.68 – 3.57 (m, 1H), 3.33 (dd, J = 12.6, 7.5 Hz, 1H), 1.88 (s, 2H). HRMS $[M+H]^+$ calcd. for $C_{18}H_{21}N_2O_4$ 329.1501, found 329.1500.

#### ***tert*-butyl (6-formylnaphthalen-2-yl) carbamate (16, an important intermediate)**

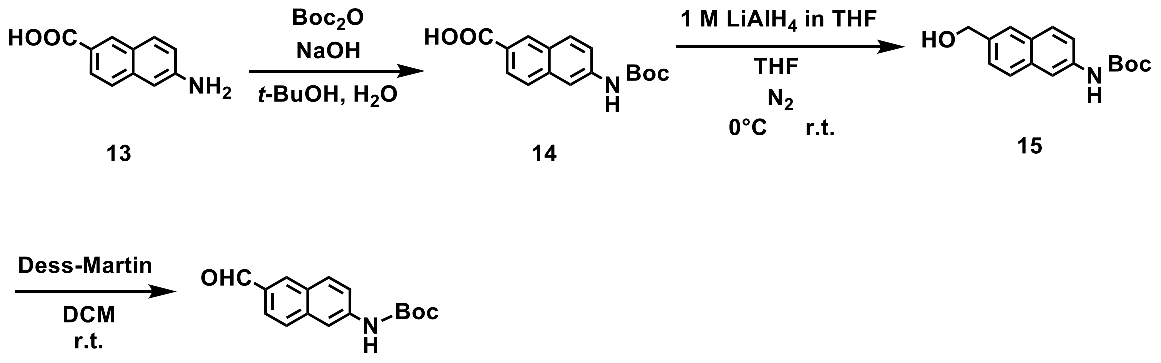

16

To a solution of **13** (4 g,1 eq) in *t*-BuOH (30 mL) and $H_2O$ (30 mL) was added $Boc_2O$ (5.9 mL, 1.2 eq) and NaOH (0.94 g, 1.1 eq). The reaction mixture was stirred at r.t, for 12 h. The reaction mixture was diluted water and washed with $CH_2Cl_2$. The aqueous layer was adjusted pH 2 with 2N HCl and extracted with ethyl acetate. The organic layer was dried over $Na_2SO_4$, and the filtrate was concentrated and purified by silica column chromatography to give **14** (5.5 g, 89%).

A stirred dispersion of $LiAlH_4$ in dry THF (48 mL) was prepared under $N_2$ and carried out at 0 °C on ice. To this solution was added dropwise the solution of **14** (2.1g, 1 eq) dissolved in dry THF (50 mL). The reaction mixture was warmed up to room temperature and subjected to stirring overnight. The progress of the reaction was monitored by TLC-analysis and once complete, the reaction mixture was quenched by slow addition of DI water at 0 °C on ice and extracted with EtOAc three times. Following which, the organic fractions were dried using $Na_2SO_4$ and concentrated *in vacuo* to result in a residue which was subjected to silica column chromatography to afford **15** (1.4 g, 80%).

To a stirred mixture of Dess-Martin (2.6 g, 1.2 eq) in dry DCM (20 mL) was added dropwise the solution of **15** (1.4 g, 1 eq) dissolved in dry DCM (40 mL). The reaction mixture was stirred at room temperature overnight. Once complete, the reaction was quenched by addition of sat. $NaHCO_3$ and sat. $Na_2S_2O_3$ and collect the organic phase. Following which, the organic fractions were dried using $Na_2SO_4$ and concentrated *in vacuo* to result in a residue which was subjected to silica column chromatography to give intermediate compound **16** (1.3 g, 92%). $^1H$ NMR (600 MHz, $CD_2Cl_2$) δ 10.08 (s, 1H), 8.26 (s, 1H), 8.09 (s, 1H), 7.94 (d, J = 8.8 Hz, 1H), 7.88 (d, J = 8.5 Hz, 1H), 7.84 (d, J = 8.5 Hz, 1H), 7.50 (d, J = 8.7 Hz, 1H), 6.99 (s, 1H), 1.55 (s, 9H).

#### 2-Amino-L-3-((6-(3-oxobut-1-en-1-yl)naphthalen-2-yl)amino)propanoic acid (AnapMe)

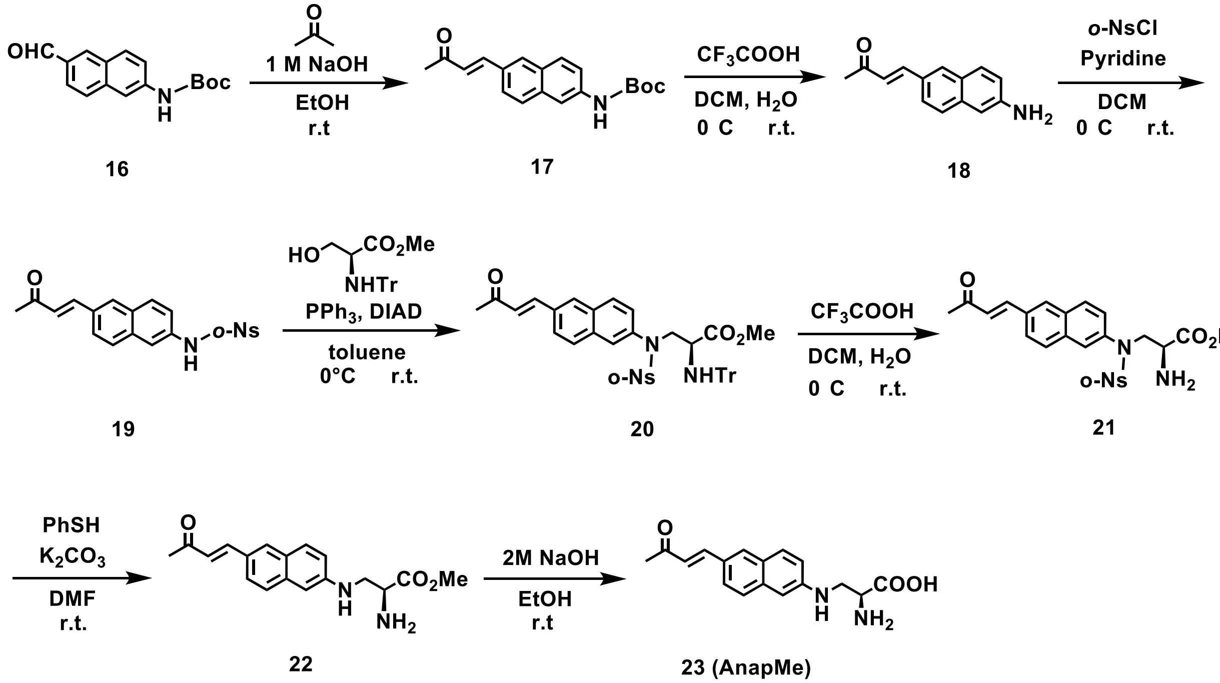

Compound **16** (300 mg, 1 eq) was mixed with EtOH (15 mL) and NaOH (1 M in $H_2O$, 1.3 mL). Then, acetone (0.2 mL, 2 eq) was added. The mixture was stirred at room temperature for 24 h. The mixture was diluted with DCM and mixed with DI H2O. The aqueous layer was extracted by DCM three times, and the combined organic layer was dried over anhydrous $Na_2SO_4$, filtered, and condensed under the reduced pressure. After evaporation, the crude solid was purified by silica chromatography obtain the desired product **17** (175 mg, 51%). $^1H$ NMR (600 MHz, CDCl3) δ 8.03 (s, 1H), 7.86 (s, 1H), 7.75 (dd, J = 11.9, 8.8 Hz, 2H), 7.64 (dd, J = 9.0, 7.4 Hz, 2H), 7.36 (dd, J = 8.8, 2.1 Hz, 1H), 6.81 – 6.76 (m, 2H), 2.41 (s, 3H), 1.56 (s, 9H). $^{13}C$ NMR (151 MHz, $CDCl_3$) δ 198.43, 152.64, 143.68, 137.39, 135.21, 130.49, 130.05, 129.64, 129.48, 128.21, 126.51, 124.17, 119.77, 114.19, 80.97, 28.32, 27.56.

To a solution of **17** (175 mg, 1 eq) in $CH_2Cl_2$ (5 mL) were added TFA (1 mL) at 0 °C. The reaction mixture was stirred at room temperature for 3 h and quenched by addition of sat. $NaHCO_3$ and collect the organic layer. The combined organic layer was washed with brine, dried over anhydrous $Na_2SO_4$, filtered, and condensed under the reduced pressure to give **18** (106 mg, 90%) without purification.

To a solution of compound **18** (105 mg, 1 eq) in $CH_2Cl_2$ (5 mL) were added pyridine (0.07 mL, 1.1 eq) and *o*-NsCl (116 mg, 1.05 eq) sequentially at 0 °C. The reaction was stirred at room temperature overnight. The reaction mixture was washed with 1 M HCl solution, water, and brine and dried over $Na_2SO_4$. The solution was concentrated, and the resulting crude product was purified through silica chromatography to give **19** (144 mg, 73%). $^1$H NMR (600 MHz, $CDCl_3$) δ 7.90 – 7.84 (m, 3H), 7.77 (d, J = 8.8 Hz, 1H), 7.75 (d, J = 8.7 Hz, 1H), 7.69 – 7.65 (m, 3H), 7.62 (d, J = 16.2 Hz, 1H), 7.59 (s, 1H), 7.53 (ddd, J = 7.8, 6.3, 1.2 Hz, 1H), 7.39 (dd, J = 8.7, 2.2 Hz, 1H), 6.81 (d, J = 16.2 Hz, 1H), 2.41 (s, 3H).

To a solution of compound **19** (144 mg, 1 eq) in toluene (10 mL) and stirred at 0 °C. To this mixture were added N-trityl-L-serine methyl ester (263 mg, 2 eq) and $PPh_3$ (1911 mg, 2 eq). DIAD (0.15 mL, 2 eq) was added dropwise to the solution. The reaction mixture was stirred overnight at room temperature. The solution was concentrated and purified by silica chromatography to give compound **20** (119 mg, 45%). $^1$H NMR (600 MHz, $CDCl_3$) δ 7.93 (s, 1H), 7.86 (d, J = 1.5 Hz, 1H), 7.79 (d, J = 8.6 Hz, 2H), 7.71 (dd, J = 8.7, 1.2 Hz, 1H), 7.67 (d, J = 16.2 Hz, 1H), 7.58 (dd, J = 7.9, 1.0 Hz, 1H), 7.55 – 7.51 (m, 1H), 7.42 – 7.39 (m, 2H), 7.43 – 7.26 (m, 7H), 7.11 – 7.03 (m, 9H), 6.85 (d, J = 16.2 Hz, 1H), 4.36 (dd, J = 14.6, 4.1 Hz, 1H), 4.30 (dd, J = 14.6, 7.0 Hz, 1H), 3.53 (dt, J = 29.0, 14.5 Hz, 1H), 3.15 (d, J = 19.5 Hz, 3H), 2.42 (s, 3H). $^{13}$C NMR (151 MHz, $CDCl_3$) δ 198.32, 173.04, 148.03, 145.54, 142.89, 137.58, 134.40, 133.92, 133.18, 132.47, 132.28, 131.82, 131.27, 130.08, 129.80, 129.15, 128.68, 128.24, 127.94, 127.90, 126.98, 126.55, 124.63, 124.08, 71.37, 56.76, 56.67, 52.06, 27.96.

To a solution of **20** (118 mg, 1 eq) in $CH_2Cl_2$ (3 mL) were added TFA (0.5 mL) and water (0.5 mL) at 0 °C. The reaction mixture was stirred at room temperature for 3 h and concentrated under vacuum. The residue was dissolved in DMF (1 mL). Thiophenol (0.04 uL, 2 eq) and $K_2CO_3$ (222 mg, 10 eq) were added to the solution sequentially. The reaction mixture was stirred at room temperature for 2 h. Water was added to the mixture, and the solution was extracted with EtOAc three times. The combined organic phase was washed with brine, dried over $Na_2SO_4$, and filtered. The filtrate was concentrated and purified by silica chromatography to give compound **22** (34 mg, 67%) as yellow solid. $^1$H NMR (600 MHz, $CDCl_3$) δ 7.79 (s, 1H), 7.67 – 7.56 (m, 4H), 6.94 (d, J = 8.5 Hz, 1H), 6.84 (s, 1H), 6.75 (d, J = 16.1 Hz, 1H), 3.84 – 3.80 (m, 1H), 3.78 (s, 3H), 3.64 (dd, J = 12.5, 3.9 Hz, 1H), 3.35 (dd, J = 12.3, 7.7 Hz, 1H), 2.40 (s, 3H). $^{13}$C NMR (151 MHz, $CDCl_3$) δ 198.50, 174.68, 146.93, 144.25, 136.46, 130.49, 129.88, 128.14, 127.17, 126.70, 125.25, 124.21, 118.68, 104.72, 53.52, 52.40, 46.89, 27.40. HRMS $[M+H]^+$ calcd. for $C_{18}H_{21}N_2O_3$ 313.1552, found 313.1549.

The solution of **22** (33 mg, 1 eq) in EtOH (3 mL) was added 2 M NaOH in water (0.5 mL). The reaction mixture was stirred at room temperature for 4 hours. The reaction mixture was then filtered and concentrated to give **23 AnapMe** as orange solid (25 mg, 80%). $^1$H NMR (600 MHz, DMSO) δ 7.95 (s, 1H), 7.76 – 7.52 (m, 6H), 7.32 (d, J = 7.7 Hz, 1H), 7.10 – 6.95 (m, 1H), 6.78 (d, J = 16.3 Hz, 2H), 3.00 (s, 2H), 2.98 (s, 1H), 2.33 (s, 3H). $^{13}$C NMR (151 MHz, DMSO) δ 198.11,

166.15, 144.37, 136.69, 130.68, 129.69, 128.64, 127.89, 127.40, 126.51, 125.76, 125.08, 124.54, 119.31, 63.31, 42.37, 27.57. HRMS $[M+H]^+$ calcd. for $C_{17}H_{19}N_2O_3$ 299.1396, found 299.1392.

#### 2-Amino-L-3-((6-(3-oxo-3-(thiophen-2-yl)prop-1-en-1-yl)naphthalen-2-yl)amino)propanoic acid (AnapTh)

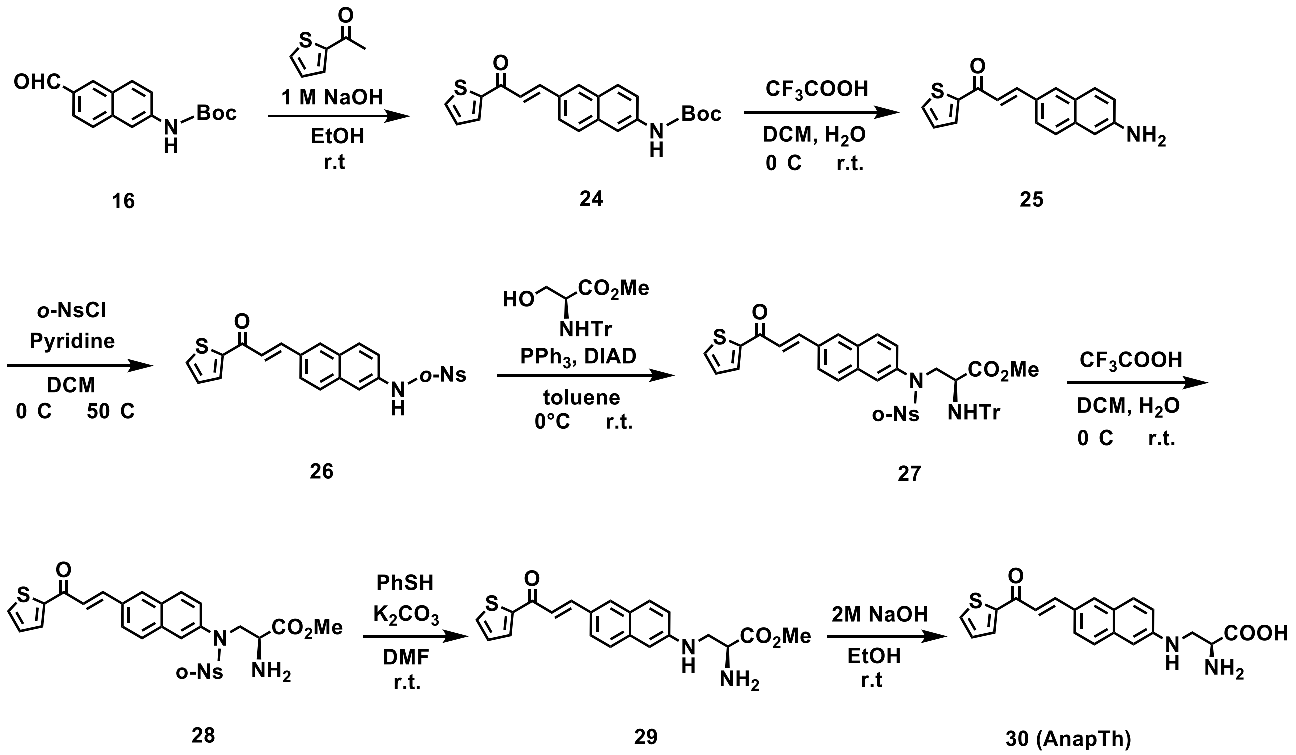

Compound **16** (300 mg, 1 eq) was mixed with EtOH (15 mL) and NaOH (1 M in $H_2O$, 1.3 mL). Then, 2-acetylthiophene (0.24 mL, 2 eq) was added. The mixture was stirred at room temperature for 24 h. The mixture was diluted with DCM and mixed with DI H2O. The aqueous layer was extracted by DCM three times, and the combined organic layer was dried over anhydrous $Na_2SO_4$, filtered, and condensed under the reduced pressure. After evaporation, the crude solid was purified by silica chromatography obtain the desired product **24** (297 mg, 70%). $^1$H NMR (600 MHz, $CDCl_3$) δ 8.07 (s, 1H), 7.98 (d, J = 15.5 Hz, 1H), 7.94 (s, 1H), 7.90 (d, J = 3.8 Hz, 1H), 7.81 – 7.74 (m, 3H), 7.69 (d, J = 4.9 Hz, 1H), 7.49 (d, J = 15.5 Hz, 1H), 7.35 (dd, J = 8.8, 2.1 Hz, 1H), 7.20 (t, J = 4.3 Hz, 1H), 6.76 (s, 1H), 1.59 – 1.53 (m, 9H). $^{13}$C NMR (151 MHz, $CDCl_3$) δ 182.26, 152.83, 145.92, 144.54, 137.59, 135.51, 134.02, 131.92, 130.99, 130.84, 129.90, 129.84, 128.47, 128.39, 124.48, 121.08, 119.93, 114.36, 81.22, 28.54.

To a solution of **24** (287 mg, 1 eq) in $CH_2Cl_2$ (25 mL) were added TFA (5 mL) at 0 °C. The reaction mixture was stirred at room temperature for 3 h and quenched by addition of sat. $NaHCO_3$ and collect the organic layer. The combined organic layer was washed with brine, dried over anhydrous $Na_2SO_4$, filtered, and condensed under the reduced pressure to give **25** (172 mg, 81%) without purification.

To a solution of compound **25** (172 mg, 1 eq) in $CH_2Cl_2$ (6 mL) were added pyridine (0.08 mL, 1.1 eq) and *o*-NsCl (144 mg, 1.05 eq) sequentially at 0 °C. The reaction was stirred at 50 °C overnight. The reaction mixture was washed with 1 M HCl solution, water, and brine and dried over $Na_2SO_4$. The solution was concentrated, and the resulting crude product was purified through silica chromatography to give **26** (150 mg, 52%).

To a solution of compound **26** (66 mg, 1 eq) in toluene (5 mL) and stir at 0 °C. To this mixture were added N-trityl-L-serine methyl ester (102 mg, 2 eq) and $PPh_3$ (74 mg, 2 eq). DIAD (0.06 mL, 2 eq) was added dropwise to the solution. The reaction mixture was stirred overnight at room temperature. The solution was concentrated and purified by silica chromatography to give compound **27** (102 mg, 90%). $^1H$ NMR (600 MHz, $CDCl_3$) δ 7.99 (dd, J = 14.3, 5.9 Hz, 2H), 7.92 (dd, J = 6.0, 2.9 Hz, 1H), 7.88 (s, 1H), 7.83 (s, 2H), 7.81 (d, J = 8.8 Hz, 1H), 7.71 – 7.68 (m, 1H), 7.61 – 7.52 (m, 3H), 7.43 – 7.39 (m, 2H), 7.33 – 7.28 (m, 2H), 7.27 (d, J = 4.2 Hz, 5H), 7.19 (dt, J = 7.9, 4.0 Hz, 1H), 7.12 – 7.05 (m, 9H), 4.36 (dt, J = 15.6, 7.9 Hz, 1H), 4.30 (dd, J = 14.6, 7.0 Hz, 1H), 3.60 – 3.51 (m, 1H), 3.22 – 3.14 (m, 3H). $^{13}C$ NMR (151 MHz, $CDCl_3$) δ 181.78, 172.84, 171.09, 147.81, 145.40, 145.32, 143.39, 137.38, 134.26, 134.13, 133.71, 133.19, 132.32, 132.09, 131.95, 131.61, 131.08, 130.05, 129.97, 128.91, 128.68, 128.47, 128.31, 128.10, 127.85, 127.69, 126.67, 126.35, 124.51, 123.86, 122.32, 71.16, 60.32, 56.59, 56.44, 51.86.

To a solution of **27** (101 mg, 1 eq) in $CH_2Cl_2$ (3 mL) were added TFA (0.5 mL) and water (0.5 mL) at 0 °C. The reaction mixture was stirred at room temperature for 3 h and concentrated under vacuum. The residue was dissolved in DMF (1 mL). Thiophenol (0.03 uL, 2 eq) and $K_2CO_3$ (175 mg, 10 eq) were added to the solution sequentially. The reaction mixture was stirred at room temperature for 2 h. Water was added to the mixture, and the solution was extracted with EtOAc three times. The combined organic phase was washed with brine, dried over $Na_2SO_4$, and filtered. The filtrate was concentrated and purified by silica chromatography to give compound **29** (22 mg, 45%) as red solid. $^1H$ NMR (600 MHz, $CDCl_3$) δ 7.98 (d, J = 15.4 Hz, 1H), 7.91 – 7.85 (m, 2H), 7.73 – 7.65 (m, 3H), 7.63 (d, J = 8.5 Hz, 1H), 7.45 (d, J = 15.4 Hz, 1H), 7.19 (t, J = 3.9 Hz, 1H), 6.95 (d, J = 8.6 Hz, 1H), 6.86 (s, 1H), 4.66 (s, 1H), 3.82 (dd, J = 14.6, 8.0 Hz, 1H), 3.80 (s, 3H), 3.65 (dd, J = 12.5, 4.1 Hz, 1H), 3.36 (dd, J = 12.6, 7.5 Hz, 1H). $^{13}C$ NMR (151 MHz, $CDCl_3$) δ 182.12, 174.70, 147.01, 145.97, 144.88, 136.60, 133.44, 131.42, 131.05, 130.06, 128.57, 128.17, 127.29, 126.68, 124.43, 119.57, 118.71, 104.82, 53.57, 52.46, 46.91. HRMS $[M+H]^+$ calcd. for $C_{21}H_{21}N_2O_3S$ 381.1273, found 381.1272.

The solution of **29** (20 mg, 1 eq) in EtOH (2 mL) was added 2 M NaOH in water (0.3 mL). The reaction mixture was stirred at room temperature for 4 hours. The reaction mixture was then filtered and concentrated to give **30 AnapTh** as red solid (15 mg, 82%). $^1H$ NMR (600 MHz, MeOD_3mm) δ 8.55 (s, 1H), 8.14 (s, 1H), 7.92 (d, J = 9.0 Hz, 2H), 7.88 (d, J = 4.3 Hz, 1H), 7.78 (d, J = 8.5 Hz, 1H), 7.67 (dd, J = 17.9, 8.8 Hz, 3H), 7.27 (d, J = 3.7 Hz, 1H), 7.02 (d, J = 8.7 Hz, 1H), 6.95 (s, 1H), 3.59 (dd, J = 14.5, 9.2 Hz, 3H), 3.35 (s, 2H). $^{13}C$ NMR (151 MHz, MeOD_3mm) δ 181.02, 170.65, 161.78, 150.01, 147.44, 147.03, 139.01, 135.81, 134.13, 132.84, 131.14, 129.97, 129.41, 128.57, 128.05, 125.50, 120.26, 105.01, 64.69, 56.86. HRMS $[M+H]^+$ calcd. for $C_{20}H_{19}N_2O_3S$ 367.1116, found 367.1115.

#### Tables

##### Table S1. Polarity of different solvents

| Solvent | Dielectric constant ε |
|---|---|
| 1,4-Dioxane | 2.25 |
| Ethyl acetate | 6.02 |
| Dichloromethane | 8.93 |
| *t*-Butanol | 10.90 |
| Isopropanol | 17.90 |
| Acetone | 20.70 |
| Ethanol | 24.50 |
| Methanol | 32.70 |
| Acetonitrile | 37.50 |
| Dimethyl sulfone | 46.70 |
| water | 80.10 |

##### Table S2. Solvent composition and viscosity of water/glycerol and ethylene glycol/glycerol mixture.

| Ratio of glycerol in water/glycerol mixture (*vol%)* | Viscosity (mPa·s) |
|---|---|
| 0% | 0.89 |
| 10% | 1.22 |
| 50% | 6.86 |
| 60% | 12.76 |
| 70% | 26.85 |
| 80% | 66.65 |
| 82% | 80.40 |
| Ratio of glycerol in ethyl glycol/glycerol mixture (*vol%)* | Viscosity (mPa·s) |

| | |
|---|---|
| 30 | 81 |
| 50 | 183 |
| 60 | 283 |
| 70 | 426 |
| 80 | 621 |

#### Table S3. List of oligonucleotide primers in this study

| Name | Aim | Sequence (5’-3’) |
|---|---|---|
| da818 | pcDNA3.1-AnapRS | GCTAGCGTTTAAACTTAAGCTTGCCACCATGGAAGAGCAATACCG |
| da821 | | GGTTTAAACGGGCCCTCTAGACTCGAGTTAAACGGGCCCGCCAAC |
| mz306 | AnapRS_W212M | CCTGACACCGTTAAAACCATGCAG |
| mz307 | | TTTAACGGTGTCAGGCATGTGATCCAGTTTATCCAGATCGTTGAG |
| mz249 | AnapRS_L500M | GCGCGCTACACTTGCCC |
| mz250 | | AAGTGTAGCGCGCCATAACCCAGGAGGACTCCATAAAGGT |
| mz249 | AnapRS_L500V | GCGCGCTACACTTGCCC |
| mz251 | | AAGTGTAGCGCGCCTCAACCCAGGAGGACTCCATAAAGGT |
| mz210 | AnapRS_L522M | CCGGTGGATATCGCGATTGGTG |
| mz211 | | CAATCGCGATATCCACCGGCATCCAGTAGTTAGCCGCTTCGG |
| mz260 | AnapRS_V524A | GATATCGCGATTGGTGGTATTGAACACG |
| mz261 | | CCACCAATCGCGATATCCGCCGGCAGCCAGTAGTTAGC |
| mz731 | AnapRS_D525N | CCGGTGAATATCGCGATTGGTGGTATTGAACACGC |
| mz732 | | TCGCGATATTCACCGGCAGCCAGTAGTTAGC |
| mz277 | AnapRS_C541A | CGCTTCTTCCACAAACTGATGCG |
| mz278 | | GTGGAAGAAGCGAGCGTAGAGACTCTCCATAATGGCGTGT |
| mz290 | AnapRS_P559G | GTTAAACAGTTGCTGTGTCAGGGTATG |
| mz291 | | AGCAACTGTTTAACTCCTTCGTCAGAGTTCACCATGCCTG |
| mz733 | AnapRS_P559A | GACGAAGCAGTTAAACAGTTGCTGTGTCAGGGTATGGTG |
| mz734 | | CTGTTTAACTGCTTCGTCAGAGTTCACCATGCCTGC |
| mz292 | AnapRS_V560G | AAACAGTTGCTGTGTCAGGGTATGGTG |
| mz293 | | GACACAGCAACTGTTTACCTGGTTCGTCAGAGTTCACCAT |
| mz216 | AnapRS_V560A | taaacagttgctgtgtcagggta |
| mz217 | | CACAGCAACTGTTTAGCTGGTTCGTCAGAGTTCACC |
| mz297 | AnapRS_L563M | CTGTGTCAGGGTATGGTGCTGG |
| mz298 | | ACCCTGACACAGCATCTGTTTAACTGGTTCGTCAGAGTTCAC |
| mz220 | AnapRS_K561Q | CAGTTGCTGTGTCAGGGTATGGT |

| | | |
|---|---|---|
| mz221 | | CCCTGACACAGCAACTGTTGAACTGGTTCGTCAGAGTTCAC |
| mz262 | AnapRS_K561R | CAGTTGCTGTGTCAGGGTATGGT |
| mz263 | | CTGACACAGCAACTGTCGAACTGGTTCGTCAGAGTTCACCA |
| mz493 | Htt-97Q | ATGGCGACCCTGGAAAAGCT |
| mz494 | | GGATCCAGGTCGGTGCAGAG |
| mz495 | pHTN for Htt-97Q | CACCGACCTGGATCCTAACTAGCATAACCCCTTGGCCGC |
| mz497 | | TTCCAGGGTCGCCATGGTGGCTTTGCTAGCCCTATAGT |
| mz503 | Htt-97Q_T3TAG | CTGGAAAAGCTGATGAAGGCCTTCG |
| mz504 | | TTCATCAGCTTTTCCAGCTACGCCATGGTGGCTTTGCTAG |
| mz505 | Htt-97Q_L7TAG | ATGAAGGCCTTCGAGTCCCTCA |
| mz506 | | ACTCGAAGGCCTTCATCTACTTTTCCAGGGTCGCCATGGT |
| mz509 | Htt-97Q_F11TAG | GCCTAGGAGTCCCTCAAAAGCTTCCAAC |
| mz510 | | CTTTTGAGGGACTCCTAGGCCTTCATCAGCTTTTCCAGGG |
| mz498 | hSOD1-A4V | ACCATGGCGACGAAGGTCGTGTGCGTGCTGAAGGG |
| mz499 | | GACCGGTGGATCCCGGG |
| mz500 | pHTN for hSOD1-A4V | CGGGATCCACCGGTCTAACTAGCATAACCCCTTGGCCGC |
| mz502 | | CGGGATCCACCGGTCATGGCAGAAATCGGTACTGGCTT |
| mz521 | hSOD1-A4V _W32TAG | TGAAGGTGTAGGGAAGCATTAAAGGACTGACTGAAGGC |
| mz522 | | AATGCTTCCCTACACCTTCACTGGTCCATTACTTTCCT |
| mz523 | hSOD1-A4V _D92TAG | TAGGGTGTGGCCGATGTGTCTATTG |
| mz524 | | ATCGGCCACACCCTATTTGTCAGCAGTCACATTGCCC |
| mz525 | hSOD1-A4V _H110TAG | AGGAGACTAGTGCATCATTGGCCGCACACT |
| mz526 | | AATGATGCACTAGTCTCCTGAGAGTGAGATCACAGAATC |
| mz549 | hSOD1-A4V _V31TAG | GAAAGTAATGGACCAGTGAAGTAGTGGGGAAGCATTAAAGGACTGACTG |
| mz550 | | CTTCACTGGTCCATTACTTTCCTTCTGC |
| mz551 | hSOD1-A4V _F45TAG | CATGTTCATGAGTTTGGAGATAATACAGCAGG |
| mz552 | | ATCTCCAAACTCATGAACATGCTATCCATGCAGGCCTTCAGTCA |
| mz553 | hSOD1-A4V _L117TAG | GTGGTCCATGAAAAAGCAGATGACTTGG |
| mz554 | | TGCTTTTTCATGGACCACCTATGTGCGGCCAATGATGCAA |
| mz707 | tagBFP | AGCGAGCTGATTAAGGAGAACATGCA |
| mz708 | | ATTAAGCTTGTGCCCCAGTTTGCTAGG |
| mz709 | pCMV6-XL5-STIM1 | GGGGCACAAGCTTAATGGCGCCAACTCTGAGGAGT |
| mz710 | | CTCCTTAATCAGCTCGCTGGCGCCCGAGCTGGT |
| mz723 | STIM1_E49TAG | gatgtggaaTAGagtgatgagttcctgagggaagacct |
| mz724 | | ctcatcactCTAttccacatccacatcaccattggc |
| mz725 | STIM1_P127TAG | gccatgTAGaggctggctgtcaccaacac |
| mz726 | | ccagcctCTAcatggcatggccactgagct |
| mz682 | STIM1_H61TAG | ctcaattaccatgacccaacagtgaaacacagcacct |
| mz683 | | tgggtcCTAgtaattgaggtcttccctcaggaactcatcac |
| mz711 | tagBFP | AGCTTGCCACCATGAGCGAGCTGATTAAGGAGAACATGCA |

| | | |
|---|---|---|
| mz712 | | CGGTGGTCCGCTACCATTAAGCTTGTGCCCCAGTTTGCT |
| mz662 | CRY2clust | AACTTAAGCTTGCCACCATGAAGATGGACAAAAAGACCATCGTCTGGT |
| mz663 | | GCCCTCTAGACTCGAGttaGCCTCCCCCTCCGCCA |
| mz641 | pcDNA3.1-tRNA | taaCTCGAGTCTAGAGGGCCCGTTTAAAC |
| mz642 | | GGTGGCAAGCTTAAGTTTAAACGCTAGCC |
| mz670 | CRY2clust_K219TAG | TAATGCCGACTAGCTGCTTAACGAGTTCATCGAAAAACAACTGATTGA |
| mz671 | | GTTAAGCAGCTAGTCGGCATTAGACCAGCCGG |
| mz674 | CRY2clust_Q263TAG | CCATGTATTTTAGTGCGCTCGGATGAAACAGATTATCTGGG |
| mz675 | | CCGAGCGCACTAAAATACATGGCGGACAGAAATCTCGCC |
| mz678 | CRY2clust_P435TAG | ACAGTGGCTCTAGGAGCTGGCCCGACTTCCTACG |
| mz679 | | CCAGCTCCTAGAGCCACTGTCGAATATACTCTCCTTCGG |

#### Table S4. Gene sequences used in this study

| Gene Name | Sequence (5'-3') |
|---|---|
| AnapRS | ATGGAAGAGCAATACCGCCCGGAAGAGATAGAATCCAAAGTACAGCTTCATTGGGATGAGAA<br>GCGCACATTTGAAGTAACCGAAGACGAGAGCAAAGAGAAGTATTACTGCTTTTCTGGCCCTC<br>CCTATCCTTCTGGTCGACTACACATGGGCCACGTACGTAACTACACCATCGGTGACGTGATCG<br>CCCGCTACCAGCGTATGCTGGGCAAAAACGTCCTGCAGCCGATCGGCTGGGACGCGTTTGG<br>TCTGCCTGCGGAAGGCGCGGCGGTGAAAAACAACACCGCTCCGGCACCGTGGACGTACGA<br>CAACATCGCGTATATGAAAAACCAGCTCAAAATGCTGGGCTTTGGTTATGACTGGAGCCGCGA<br>GCTGGCAACCTGTACGCCGGAATACTACCGTTGGGAACAGAAATTCTTCACCGAGCTGTATAA<br>AAAAGGCCTGGTATATAAGAAGACTTCTGCGGTCAACTGGTGTCCGAACGACCAGACCGTAC<br>TGGCGAACGAACAAGTTATCGACGGCTGCTGCTGGCGCTGCGATACCAAAGTTGAACGTAAA<br>GAGATCCCGCAGTGGTTTATCAAAATCACTGCTTACGCTGACGAGCTGCTCAACGATCTGGAT<br>AAACTGGATCACTGGCCTGACACCGTTAAAACCATGCAGCGTAACTGGATCGGTCGTTCCGA<br>AGGCGTGGAGATCACCTTCAACGTTAACGACTATGACAACACGCTGACCGTTTACACTACCC<br>GCCCGGACACCTTTATGGGTTGTACCTACCTGGCGGTAGCTGCGGGTCATCCGCTGGCGCA<br>GAAAGCGGCGGAAAATAATCCTGAACTGGCGGCCTTTATTGACGAATGCCGTAACACCAAAG<br>TTGCCGAAGCTGAAATGGCGACGATGGAGAAAAAAGGCGTCGATACTGGCTTTAAAGCGGTT<br>CACCCATTAACGGGCGAAGAAATTCCCGTTTGGGCAGCAAACTTCGTATTGATGGAGTACGG<br>CACGGGCGCAGTTATGGCGGTACCGGGGCACGACCAGCGCGACTACGAGTTTGCCTCTAAA<br>TACGGCCTGAACATCAAACCGGTTATCCTGGCAGCTGACGGCTCTGAGCCAGATCTTTCTCA<br>GCAAGCCCTGACTGAAAAAGGCGTGCTGTTCAACTCTGGCGAGTTCAACGGTCTTGACCAT<br>GAAGCGGCCTTCAACGCCATCGCCGATAAACTGACTGCGATGGGCGTTGGCGAGCGTAAAG<br>TGAACTACCGCCTGCGCGACTGGGGTGTTTCCCGTCAGCGTTACTGGGGCGCGCCGATTCC<br>GATGGTGACTCTAGAAGACGGTACCGTAATGCCGACCCCGGACGACCAGCTGCCGGTGATC<br>CTGCCGGAGGATGTGGTAATGGACGGCATTACCAGCCCGATTAAAGCAGATCCGGAGTGGG<br>CGAAAACTACCGTTAACGGTATGCCAGCACTGCGTGAAACCGACACTTTCGACACCTTTATGG<br>AGTCCTCCTGGGTTCTTGCGCGCTACACTTGCCCGCAGTACAAAGAAGGTATGCTGGATTCC<br>GAAGCGGCTAACTACTGGCTGCCGGTGGATATCGCGATTGGTGGTATTGAACACGCCATTAT<br>GGAGAGTCTCTACTGTCGCTTCTTCCACAAACTGATGCGTGATGCAGGCATGGTGAACTCTG<br>ACGAACCAGTTAAACAGTTGCTGTGTCAGGGTATGGTGCTGGCAGATGCCTTCTACTATGTTG<br>GCGAAAACGGCGAACGTAACTGGGTTTCCCCGGTTGATGCTATCGTTGAACGTGACGAGAAA<br>GGCCGTATCGTGAAAGCGAAAGATGCGGCAGGCCATGAACTGGTTTATACCGGCATGAGCAA<br>AATGTCCAAGTCGAAGAACAACGGTATCGACCCGCAGGTGATGGTTGAACGTTACGGCGCG<br>GACACCGTTCGTCTGTTTATGATGTTTGCTTCTCCGGCTGATATGACTCTCGAATGGCAGGAA<br>TCCGGTGTGGAAGGGGCTAACCGCTTCCTGAAACGTGTCTGGAAACTGGTTTACGAGCACA<br>CAGCAAAAAGTGATGTTGCGGCACTGAACGTTGATGCGCTGACTGAAAATCAGAAAGCGCTG<br>CGTCGCGATGTGCATAAAACGATCGCTAAAGTGACCGATGATATCGGCCGTCGTCAGACCTT<br>CAACACCGCAATTGCGGCGATTATGGAGCTGATGAACAAACTGGCGAAAGCACCAACCGATG<br>GCGAGCAGGATCGCGCTCTGATGCAGGAAGCACTGCTGGCCGTTGTCCGTATGCTTAACCC<br>GTTCACCCCGCACATCTGCTTCACGCTGTGGCAGGAACTGAAAGGCGAAGGCGATATCGAC<br>AACGCGCCGTGGCCGGTTGCTGACGAAAAAGCGATGGTGGAAGACTCCACGCTGGTCGTG |

| | |
|---|---|
| | GTGCAGGTTAACGGTAAAGTCCGTGCCAAAATCACCGTTCCGGTGGACGCAACGGAAGAAC<br>AGGTTCGCGAACGTGCTGGCCAGGAACATCTGGTAGCAAAATATCTTGATGGCGTTACTGTA<br>CGTAAAGTGATTTACGTACCAGGTAAACTCCTCAATCTGGTCGTTGGCGGGCCCGTTTAA |
| tRNA$_{CUA}^{\mathit{Ec}\mathrm{Leu}}$ | GCCCGGATGGTGGAATCGGTAGACACAAGGGATTCTAAATCCCTCGGCGTTCGCGCTGTGC<br>GGGTTCAAGTCCCGCTCCGGGTA |
| hSOD1-A4V | ATGGCGACGAAGGTCGTGTGCGTGCTGAAGGGCGACGGCCCAGTGCAGGGCATCATCAATT<br>TCGAGCAGAAGGAAAGTAATGGACCAGTGAAGGTGTGGGGAAGCATTAAAGGACTGACTGA<br>AGGCCTGCATGGATTCCATGTTCATGAGTTTGGAGATAATACAGCAGGCTGTACCAGTGCAGG<br>TCCTCACTTTAATCCTCTATCCAGAAAACACGGTGGGCCAAAGGATGAAGAGAGGCATGTTG<br>GAGACTTGGGCAATGTGACTGCTGACAAAGATGGTGTGGCCGATGTGTCTATTGAAGATTCT<br>GTGATCTCACTCTCAGGAGACCATTGCATCATTGGCCGCACACTGGTGGTCCATGAAAAAGC<br>AGATGACTTGGGCAAAGGTGGAAATGAAGAAAGTACAAAGACAGGAAACGCTGGAAGTCGTT<br>TGGCTTGTGGTGTAATTGGGATCGCCCAAGGGTCGACGGTACCGCGGGCCCGGGATCCACC<br>GGTC |
| Htt-97Q | ATGGCGACCCTGGAAAAGCTGATGAAGGCCTTCGAGTCCCTCAAAAGCTTCCAACAGCAGC<br>AACAGCAACAACAGCAGCAACAGCAACAACAGCAGCAACAGCAACAACAGCAGCAACAGCA<br>ACAACAGCAGCAACAGCAACAACAGCAGCAACAGCAACAACAGCAGCAACAGCAACAACAG<br>CAGCAACAGCAACAACAGCAGCAACAGCAACAACAGCAGCAACAGCAACAACAGCAGCAAC<br>AGCAACAACAGCAGCAACAGCAACAACAGCAGCAACAGCAACAACAGCAGCAACAGCAACA<br>ACAGCAGCAACAGCAACAACAGCAGCAACAGCAACAACCGCCACCACCTCCCCCTCCACCC<br>CCACCTCCTCAACTTCCTCAACCTCCTCCACAGGCACAGCCTCTGCTGCCTCAGCCACAACC<br>TCCTCCACCTCCACCTCCACCTCCTCCAGGCCCAGCTGTGGCTGAGGAGCCTCTGCACCGA<br>CCTGGATCC |
| tagBFP | AGCGAGCTGATTAAGGAGAACATGCACATGAAGCTGTACATGGAGGGCACCGTGGACAACCA<br>TCACTTCAAGTGCACATCCGAGGGCGAAGGCAAGCCCTACGAGGGCACCCAGACCATGAGA<br>ATCAAGGTGGTCGAGGGCGGCCCTCTCCCCTTCGCCTTCGACATCCTGGCTACTAGCTTCCT<br>CTACGGCAGCAAGACCTTCATCAACCACACCCAGGGCATCCCCGACTTCTTCAAGCAGTCCT<br>TCCCTGAGGGCTTCACATGGGAGAGAGTCACCACATACGAAGACGGGGGCGTGCTGACCGC<br>TACCCAGGACACCAGCCTCCAGGACGGCTGCCTCATCTACAACGTCAAGATCAGAGGGGTG<br>AACTTCACATCCAACGGCCCTGTGATGCAGAAGAAAACACTCGGCTGGGAGGCCTTCACCG<br>AGACGCTGTACCCCGCTGACGGCGGCCTGGAAGGCAGAAACGACATGGCCCTGAAGCTCG<br>TGGGCGGGAGCCATCTGATCGCAAACATCAAGACCACATATAGATCCAAGAAACCCGCTAAG<br>AACCTCAAGATGCCTGGCGTCTACTATGTGGACTACAGACTGGAAAGAATCAAGGAGGCCAA<br>CAACGAGACCTACGTCGAGCAGCACGAGGTGGCAGTGGCCAGATACTGCGACCTCCCTAGC<br>AAACTGGGGCACAAGCTTAAT |
| STIM1 | AACTCTGAGGAGTCCACTGCAGCAGAGTTTTGCCGAATTGACAAGCCCCTGTGTCACAGTGA<br>GGATGAGAAACTCAGCTTCGAGGCAGTCCGTAACATCCACAAACTGATGGACGATGATGCCA<br>ATGGTGATGTGGATGTGGAAGAAAGTGATGAGTTCCTGAGGGAAGACCTCAATTACCATGAC<br>CCAACAGTGAAACACAGCACCTTCCATGGTGAGGATAAGCTCATCAGCGTGGAGGACCTGTG<br>GAAGGCATGGAAGTCATCAGAAGTATACAATTGGACCGTGGATGAGGTGGTACAGTGGCTGA<br>TCACATATGTGGAGCTGCCTCAGTATGAGGAGACCTTCCGGAAGCTGCAGCTCAGTGGCCAT<br>GCCATGCCAAGGCTGGCTGTCACCAACACCACCATGACAGGGACTGTGCTGAAGATGACAG<br>ACCGGAGTCATCGGCAGAAGCTGCAGCTGAAGGCTCTGGATACAGTGCTCTTTGGGCCTCC<br>TCTCTTGACTCGCCATAATCACCTCAAGGACTTCATGCTGGTGGTGTCTATCGTTATTGGTGT<br>GGGCGGCTGCTGGTTTGCCTATATCCAGAACCGTTACTCCAAGGAGCACATGAAGAAGATGA<br>TGAAGGACTTGGAGGGGTTACACCGAGCTGAGCAGAGTCTGCATGACCTTCAGGAAAGGCT<br>GCACAAGGCCCAGGAGGAGCACCGCACAGTGGAGGTGGAGAAGGTCCATCTGGAAAAGAA<br>GCTGCGCGATGAGATCAACCTTGCTAAGCAGGAAGCCCAGCGGCTGAAGGAGCTGCGGGA<br>GGGTACTGAGAATGAGCGGAGCCGCCAAAAATATGCTGAGGAGGAGTTGGAGCAGGTTCGG<br>GAGGCCTTGAGGAAAGCAGAGAAGGAGCTAGAATCTCACAGCTCATGGTATGCTCCAGAGG<br>CCCTTCAGAAGTGGCTGCAGCTGACACATGAGGTGGAGGTGCAATATTACAACATCAAGAAG<br>CAAAATGCTGAGAAGCAGCTGCTGGTGGCCAAGGAGGGGGCTGAGAAGATAAAAAAGAAGA<br>GAAACACACTCTTTGGCACCTTCCACGTGGCCCACAGCTCTTCCCTGGATGATGTAGATCATA<br>AAATTCTAACAGCTAAGCAAGCACTGAGCGAGGTGACAGCAGCATTGCGGGAGCGCCTGCA<br>CCGCTGGCAACAGATCGAGATCCTCTGTGGCTTCCAGATTGTCAACAACCCTGGCATCCACT<br>CACTGGTGGCTGCCCTCAACATAGACCCCAGCTGGATGGGCAGTACACGCCCCAACCCTGC<br>TCACTTCATCATGACTGACGACGTGGATGACATGGATGAGGAGATTGTGTCTCCCTTGTCCAT<br>GCAGTCCCCTAGCCTGCAGAGCAGTGTTCGGCAGCGCCTGACGGAGCCACAGCATGGCCT<br>GGGATCTCAGAGGGATTTGACCCATTCCGATTCGGAGTCCTCCCTCCACATGAGTGACCGCC<br>AGCGTGTGGCCCCCAAACCTCCTCAGATGAGCCGTGCTGCAGACGAGGCTCTCAATGCCAT |

| | |
|---|---|
| | GACTTCCAATGGCAGCCACCGGCTGATCGAGGGGGTCCACCCAGGGTCTCTGGTGGAGAA<br>ACTGCCTGACAGCCCTGCCCTGGCCAAGAAGGCATTACTGGCGCTGAACCATGGGCTGGAC<br>AAGGCCCACAGCCTGATGGAGCTGAGCCCCTCAGCCCCACCTGGTGGCTCTCCACATTTGG<br>ATTCTTCCCGTTCTCACAGCCCCAGCTCCCCAGACCCAGACACACCATCTCCAGTTGGGGAC<br>AGCCGAGCCCTGCAAGCCAGCCGAAACACACGCATTCCCCACCTGGCTGGCAAGAAGGCT<br>GTGGCTGAGGAGGATAATGGCTCTATTGGCGAGGAAACAGACTCCAGCCCAGGCCGGAAGA<br>AGTTTCCTCTCAAAATCTTTAAGAAGCCTCTTAAGAAG |
| CRY2clust (with 3xS7G tale) | AAGATGGACAAAAAGACCATCGTCTGGTTTCGGAGAGATTTGAGAATAGAAGATAATCCCGCG<br>CTCGCCGCCGCGGCCCACGAGGGTTCCGTCTTCCCCGTTTTCATTTGGTGTCCTGAAGAAG<br>AAGGCCAGTTTTATCCCGGAAGGGCCTCTAGGTGGTGGATGAAGCAAAGTCTGGCCCATCTT<br>AGCCAGTCACTGAAAGCACTGGGCAGTGATCTTACCCTGATCAAGACACACAATACCATCTCT<br>GCCATCCTCGACTGCATCAGGGTGACCGGCGCAACGAAAGTCGTGTTTAACCACCTGTACGA<br>TCCAGTTAGTCTGGTGCGCGACCACACTGTGAAGGAGAAGCTGGTGGAACGGGGGATCAGT<br>GTGCAGAGCTACAACGGGGACCTTCTGTACGAGCCATGGGAGATCTATTGCGAGAAAGGGA<br>AACCGTTCACCTCCTTCAACAGTTACTGGAAGAAATGTTTGGATATGTCAATAGAGTCCGTTAT<br>GTTGCCCCCTCCCTGGAGACTGATGCCGATTACTGCTGCTGCAGAGGCCATCTGGGCCTGC<br>TCCATCGAGGAACTCGGTCTGGAAAATGAAGCAGAAAAGCCAAGCAATGCACTTCTCACTAG<br>AGCCTGGAGCCCCGGCTGGTCTAATGCCGACAAGCTGCTTAACGAGTTCATCGAAAAACAAC<br>TGATTGACTACGCGAAGAACTCCAAGAAAGTGGTAGGTAACTCAACTAGCTTGCTCTCTCCAT<br>ATCTCCATTTTGGCGAGATTTCTGTCCGCCATGTATTTCAGTGCGCTCGGATGAAACAGATTAT<br>CTGGGCTCGCGATAAAAACAGCGAAGGCGAAGAAAGCGCCGATCTGTTCcTGCGAGGGATC<br>GGACTTCGGGAATACTCCCGGTATATATGTTTCAACTTTCCATTCACACACGAGCAGAGTCTG<br>TTGTCCCACCTCAGGTTCTTCCCCTGGGACGCCGATGTCGACAAATTCAAGGCATGGAGACA<br>GGGAAGGACAGGGTACCCACTCGTGGATGCTGGCATGAGAGAGCTCTGGGCTACAGGCTG<br>GATGCACAACCGCATCCGGGTAATCGTGTCCTCATTTGCTGTCAAGTTTCTGCTCCTGCCTTG<br>GAAATGGGGAATGAAGTACTTTTGGGATACCCTTCTCGACGCCGACTTGGAGTGTGACATTCT<br>GGGATGGCAATATATTAGCGGGTCAATTCCTGACGGCCATGAGTTGGACAGGTTGGACAATC<br>CGGCCTTGCAGGGAGCTAAGTATGATCCCGAAGGAGAGTATATTCGACAGTGGCTCCCCGAG<br>CTGGCCCGACTTCCTACGGAGTGGATTCACCATCCTTGGGACGCACCACTGACAGTGCTCAA<br>GGCAAGCGGGGTGGAGCTGGGCACCAATTACGCTAAGCCTATAGTTGATATAGATACAGCAC<br>GCGAGCTGCTGGCTAAAGCGATCTCTCGCACTCGGGAGGCGCAGATTATGATCGGTGCTGC<br>CGCAAGGGACCCTCCTGATCTTGACAATGTGTACAGTGGGGGAGGCGGAGGGGGTGGTTCA<br>GGTGGTGGTGGTGGTGGTGGTTCGGGTGGTGGCGGAGGGGGAGGC |

#### Supplementary figures

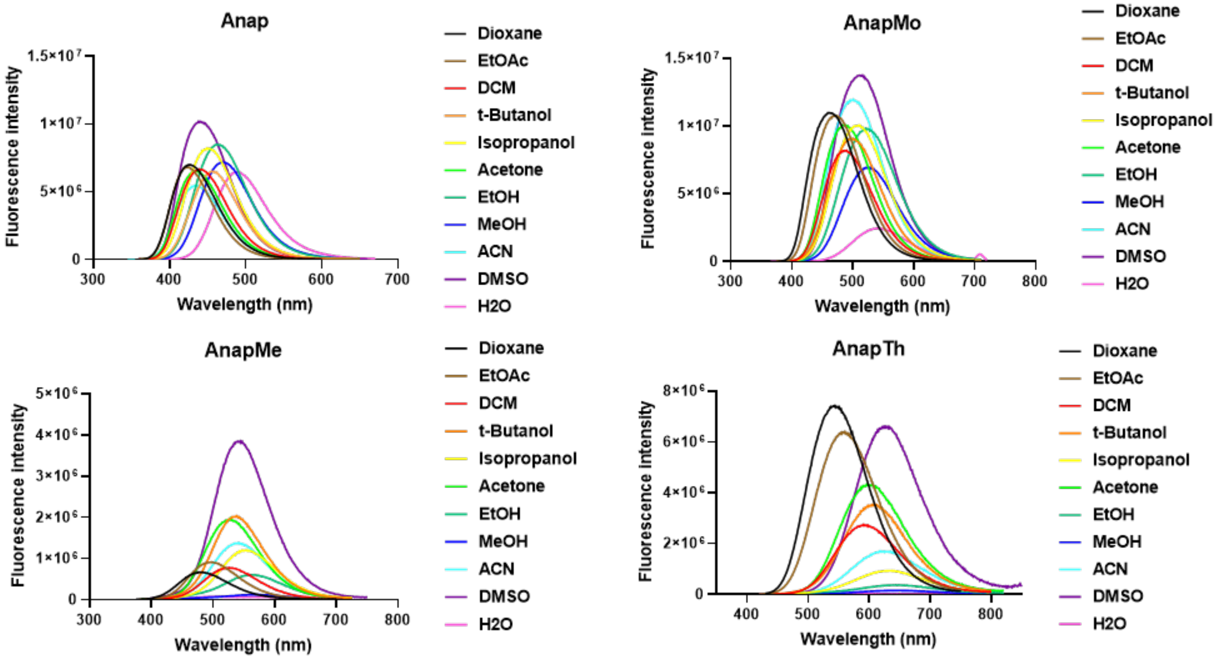

**Figure S1. Fluorescence spectra of Anap, Anap-methoxy, Anap-methyl and Anap-thiophene in different solvents.**

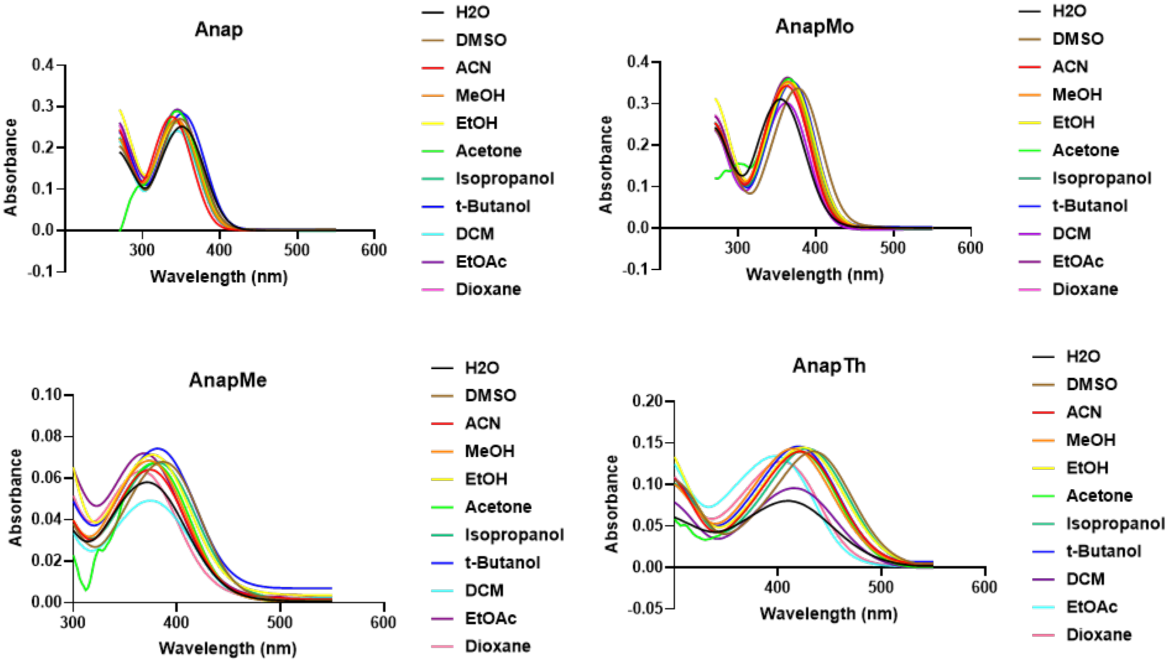

**Figure S2. Absorption spectra of Anap, AnapMo, AnapMe and AnapTh in different solvents.**

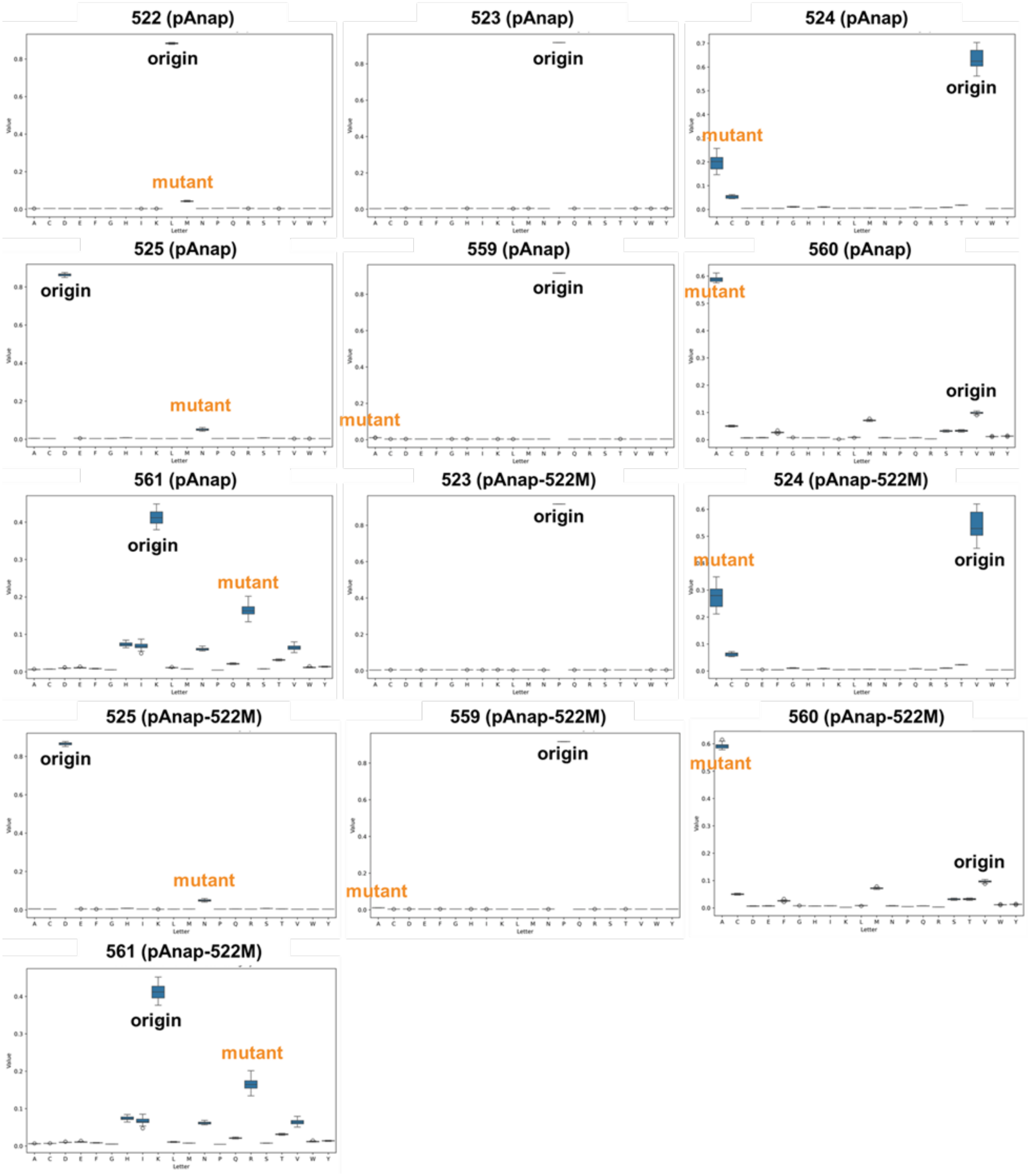

**Figure S3. Value distribution scores of each natural amino acid for the conflicted residues in AnapRS active pocket.**

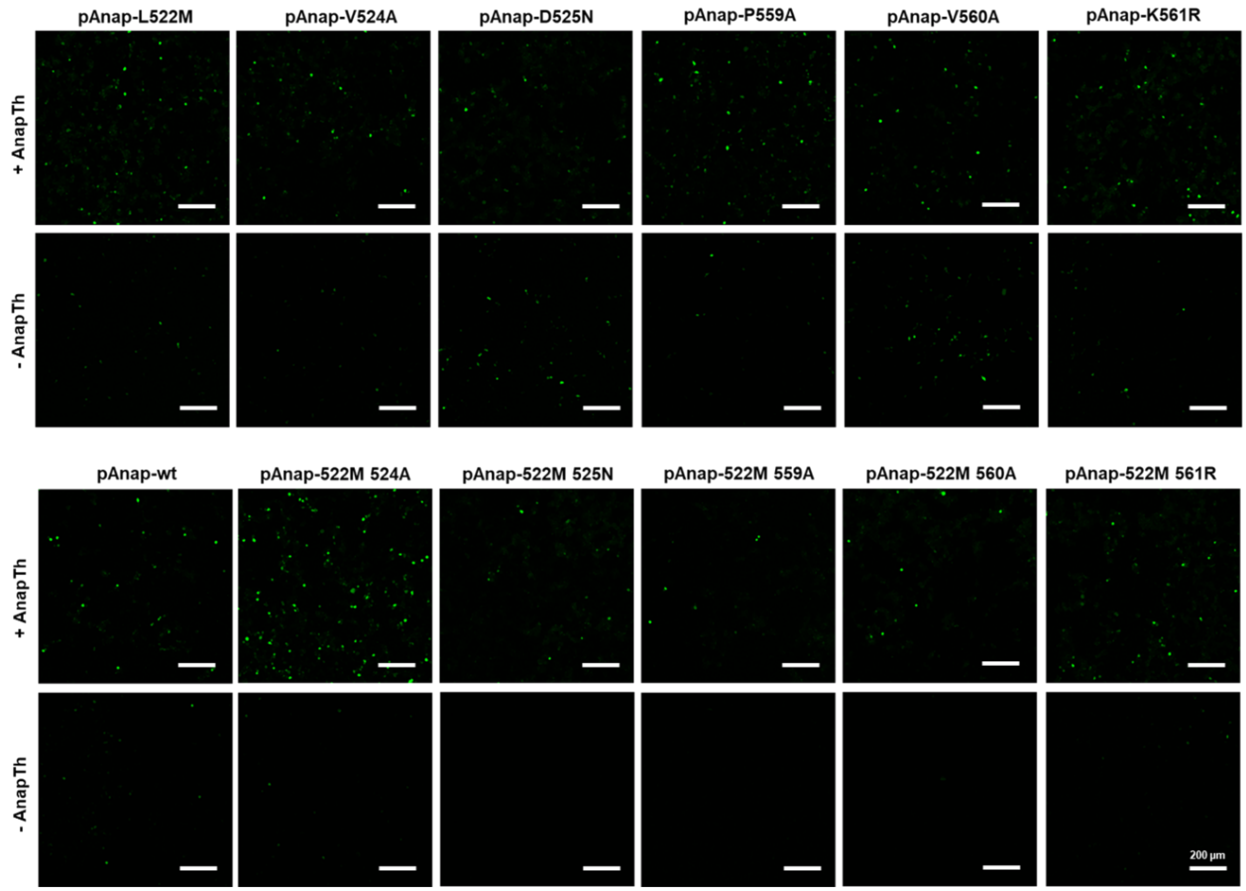

**Figure S4. Confocal images of HEK293T cells expressing EGFP-39AnapTh using Anap RS mutations.** Scale = 200 μm**.**

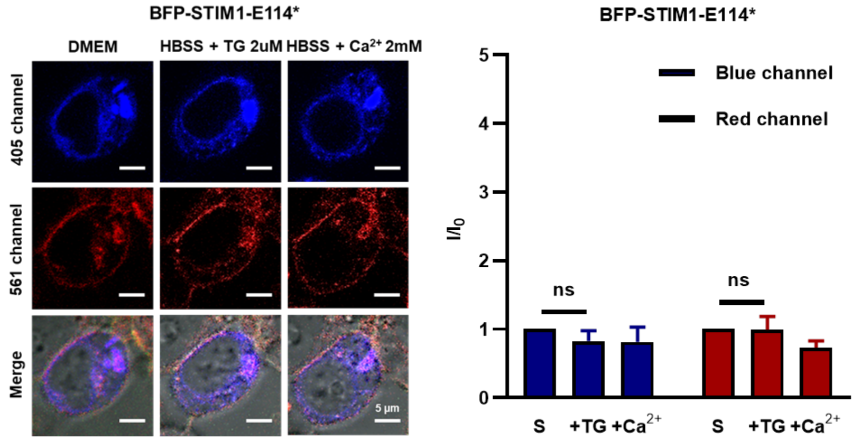

**Figure S5. Confocal imaging and fluorescence changes of HEK293T cells expressing BFP-STIM1-E114AnapTh.** DMEM medium was firstly replaced with HBSS ($Ca^{2+}$ free) including 2 μM TG. After 15 minutes, the TG included HBSS was replaced with HBSS with 2 mM $CaCl_2$. The fluorescence of BFP and AnapTh was collected separately via 405 nm blue channel and 561 nm red channel.

#### NMR

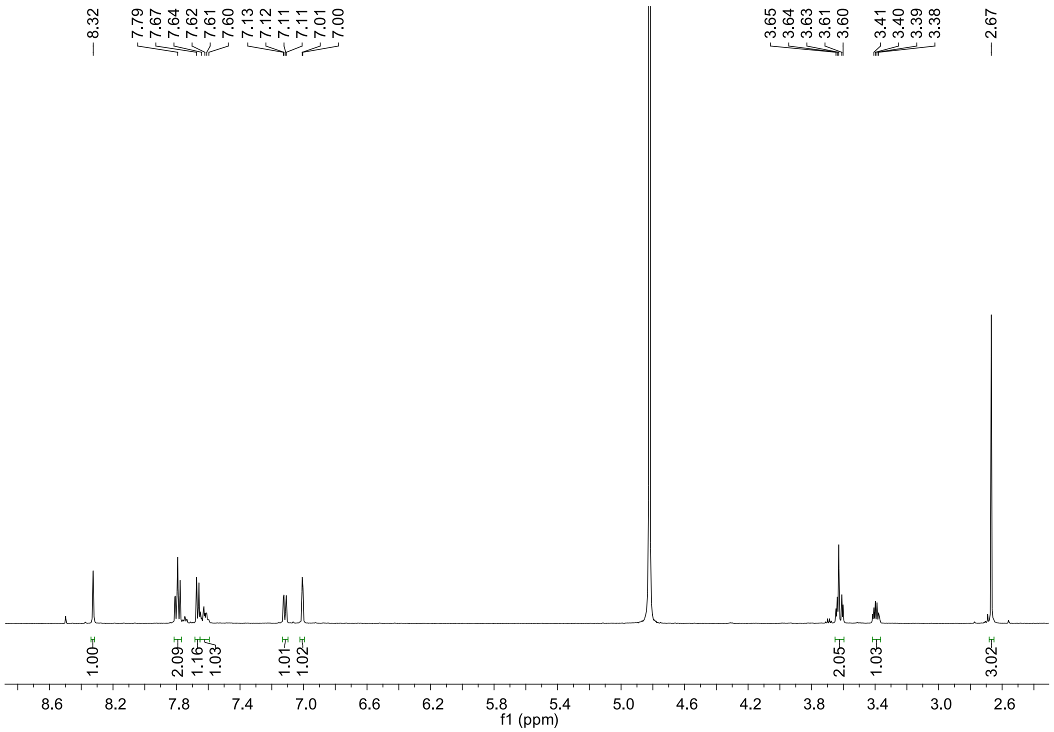

Figure S6. $^1$H NMR of Anap.

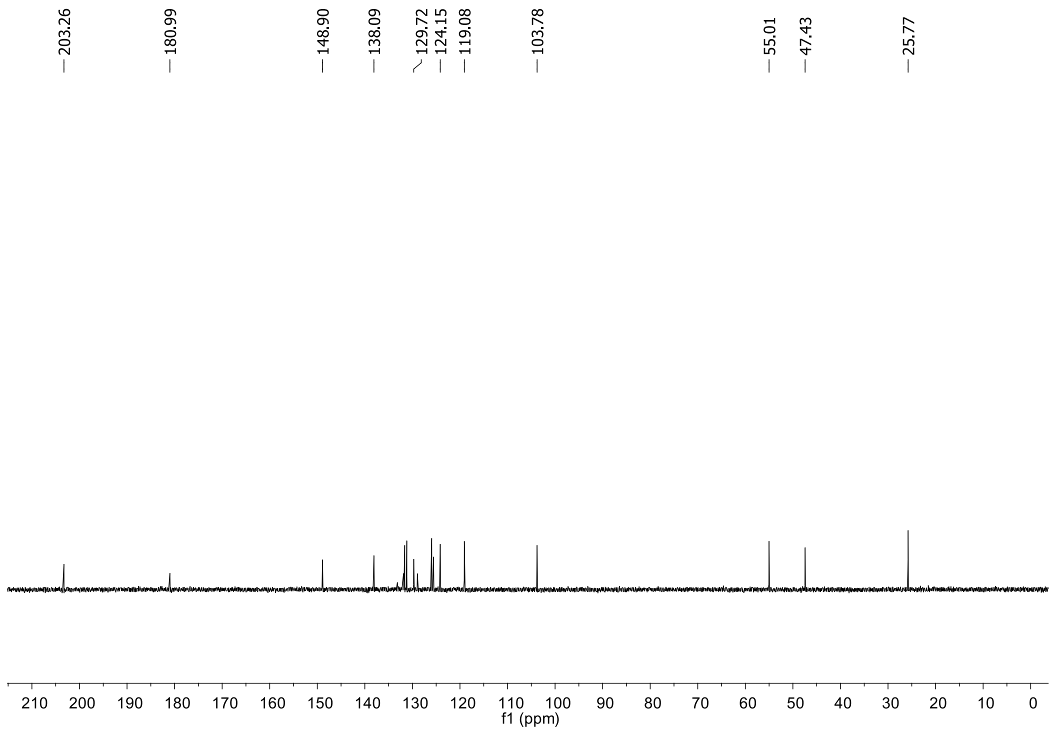

Figure S7. $^{13}$C NMR of L-Anap.

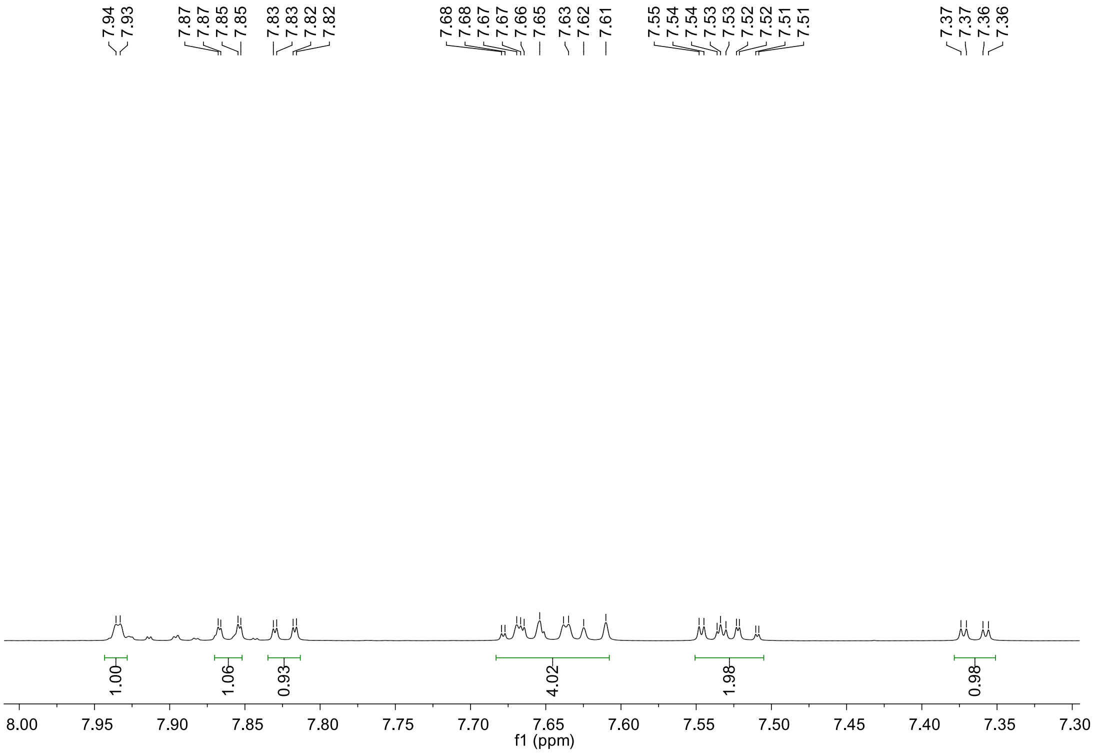

Figure S8. [1]H NMR of compound **8**.

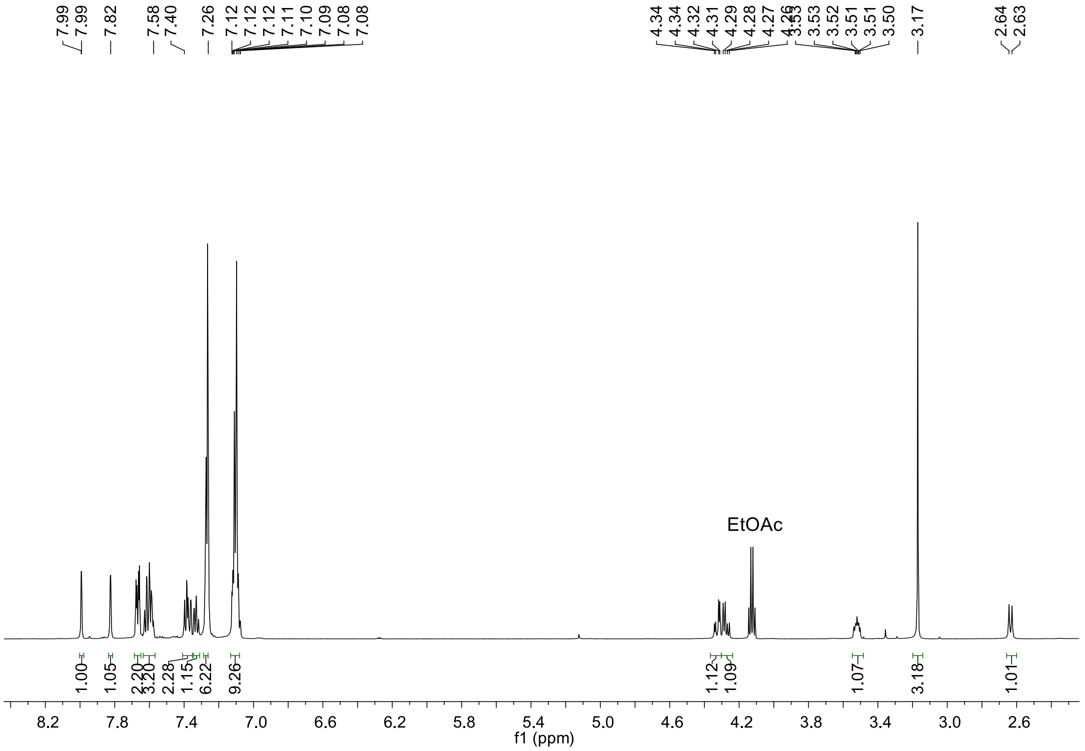

Figure S9. [1]H NMR of compound **9**.

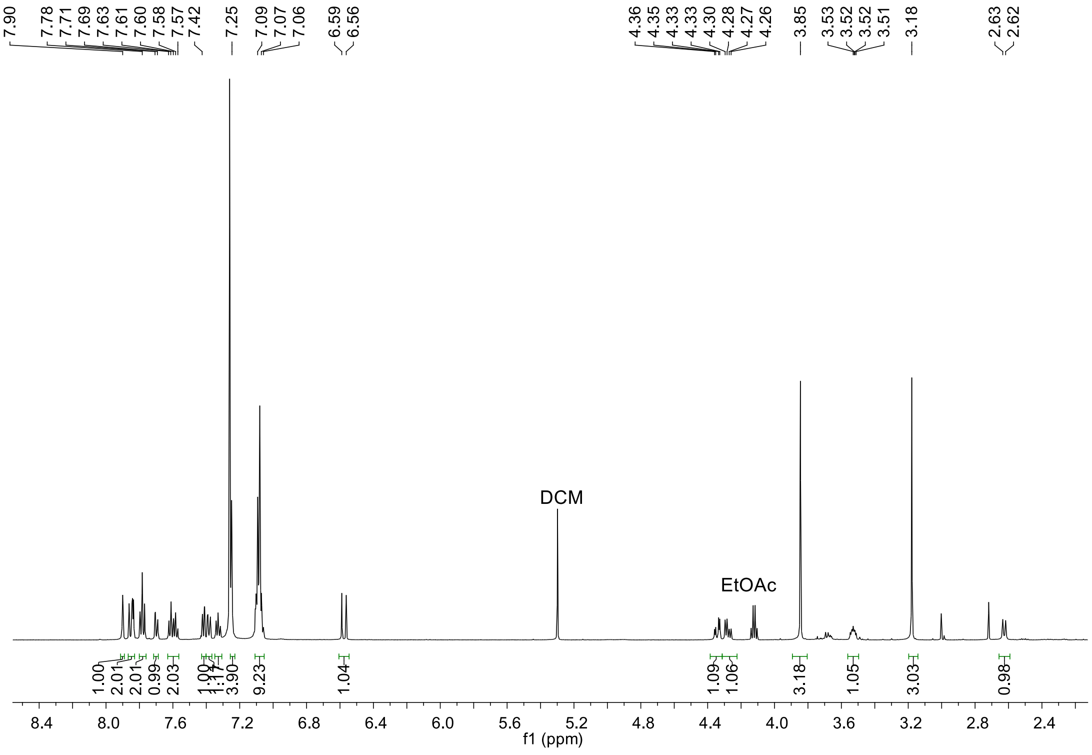

Figure S10. $^{1}$H NMR of compound **10**.

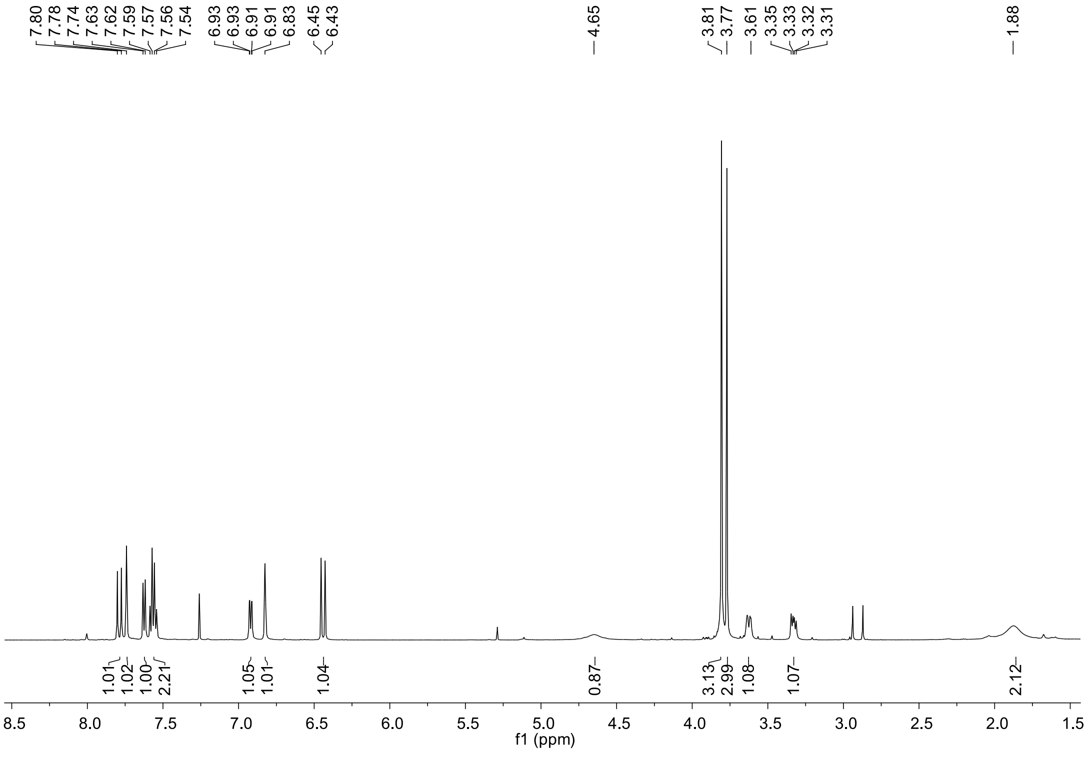

Figure S11. $^{1}$H NMR of compound **12**.

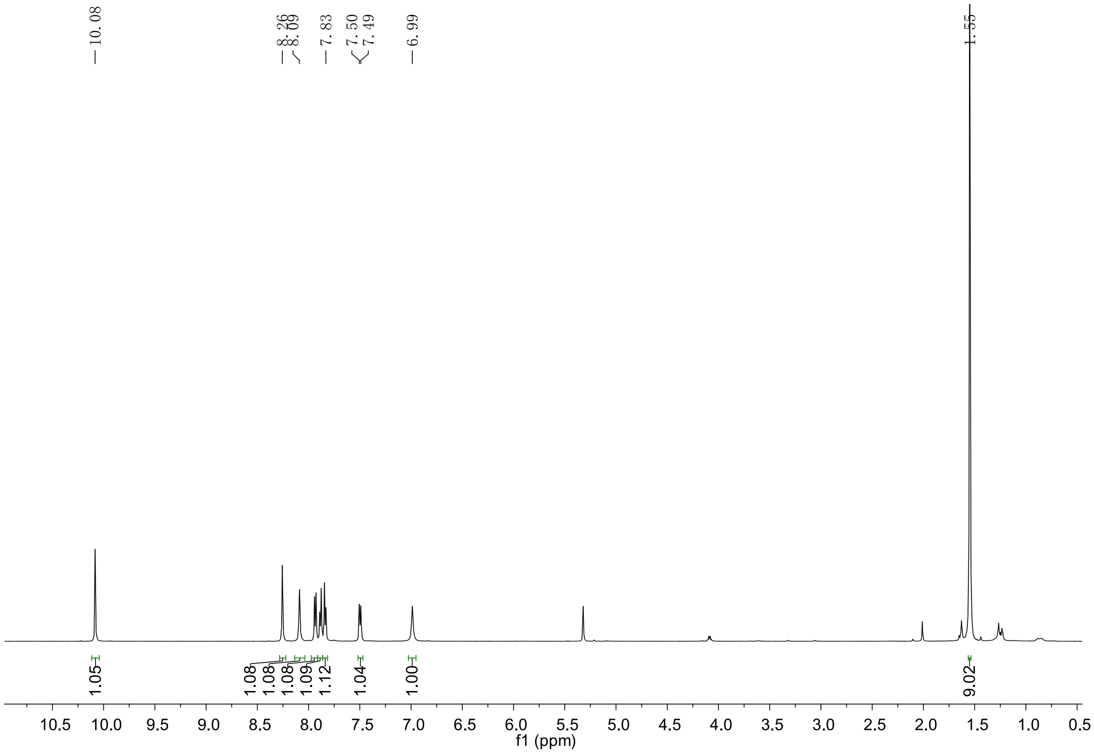

Figure S12. $^{1}$H NMR of compound **16**.

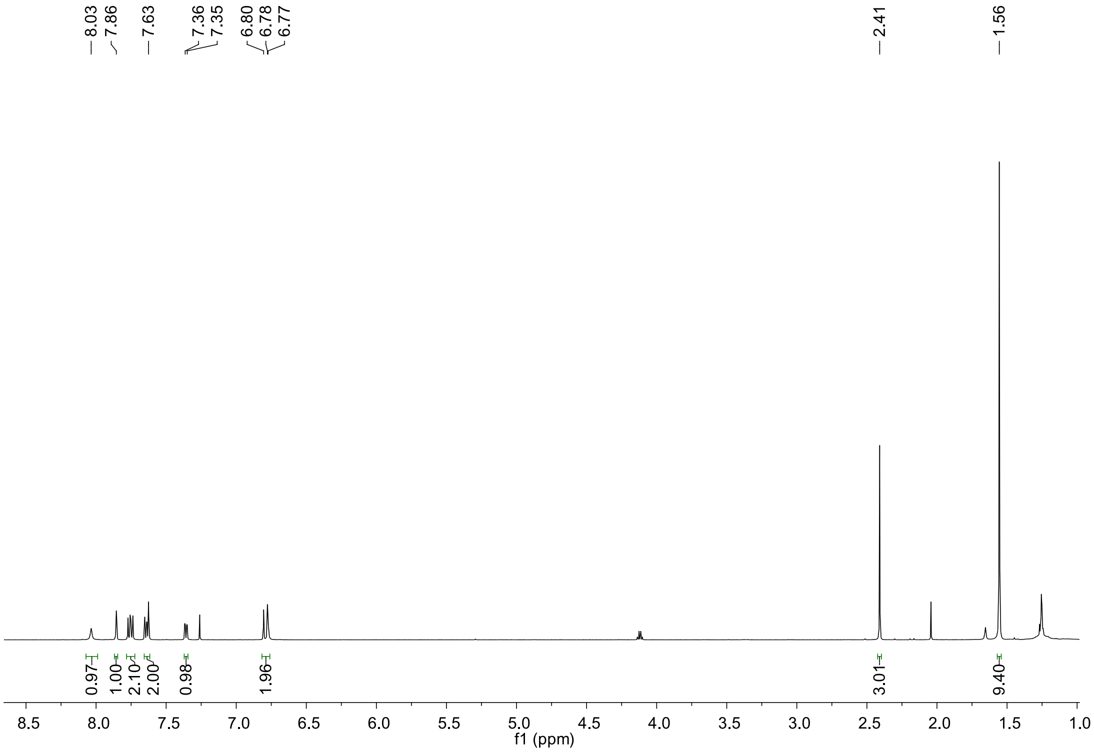

Figure S13. $^{1}$H NMR of compound **17**.

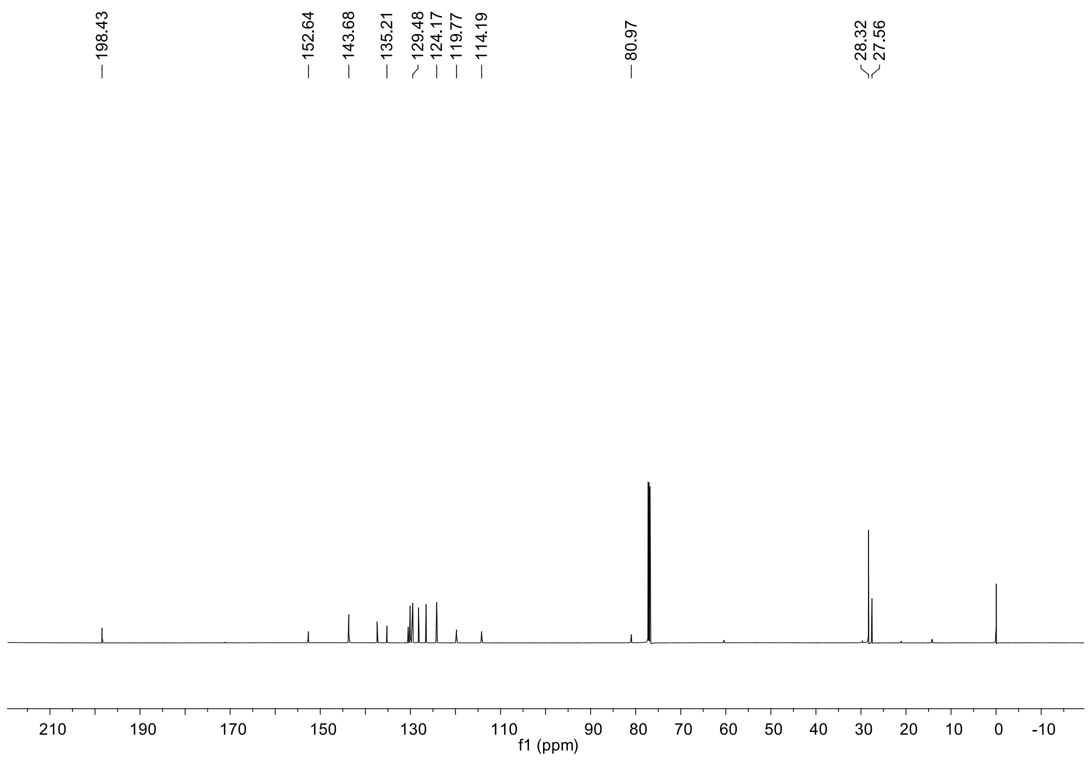

Figure S14. $^{13}C$ NMR of compound **17**.

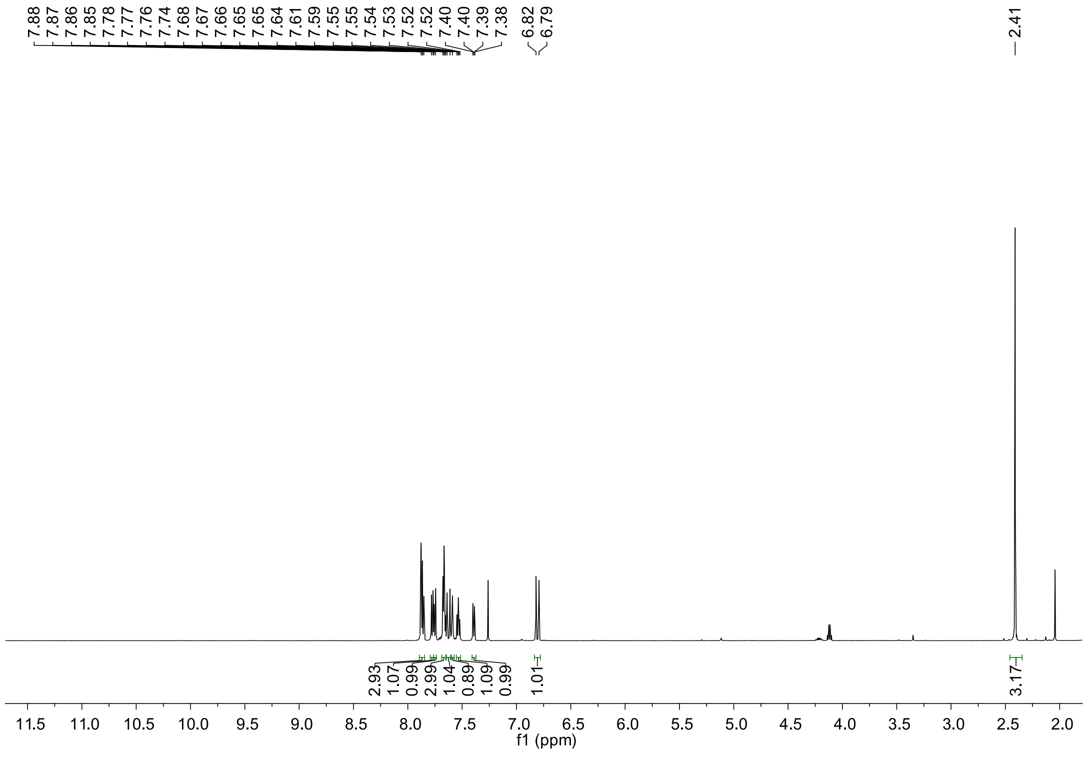

Figure S15. $^{1}H$ NMR of compound **19**.

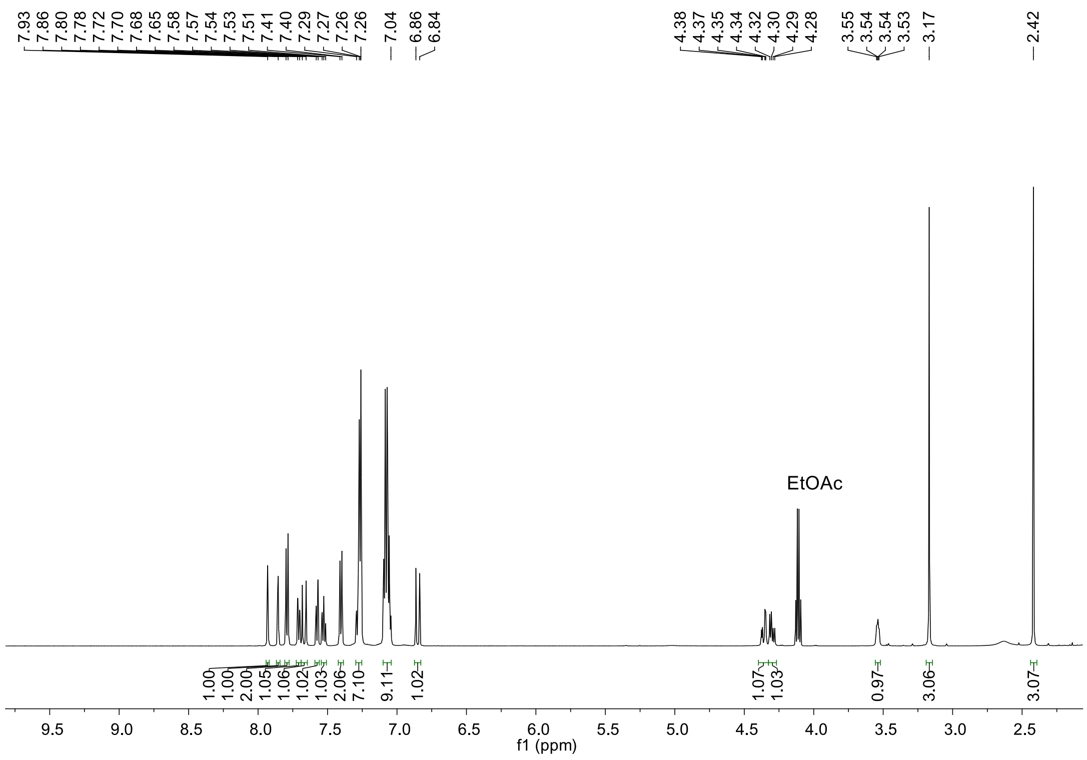

Figure S16. $^{1}$H NMR of compound **20**.

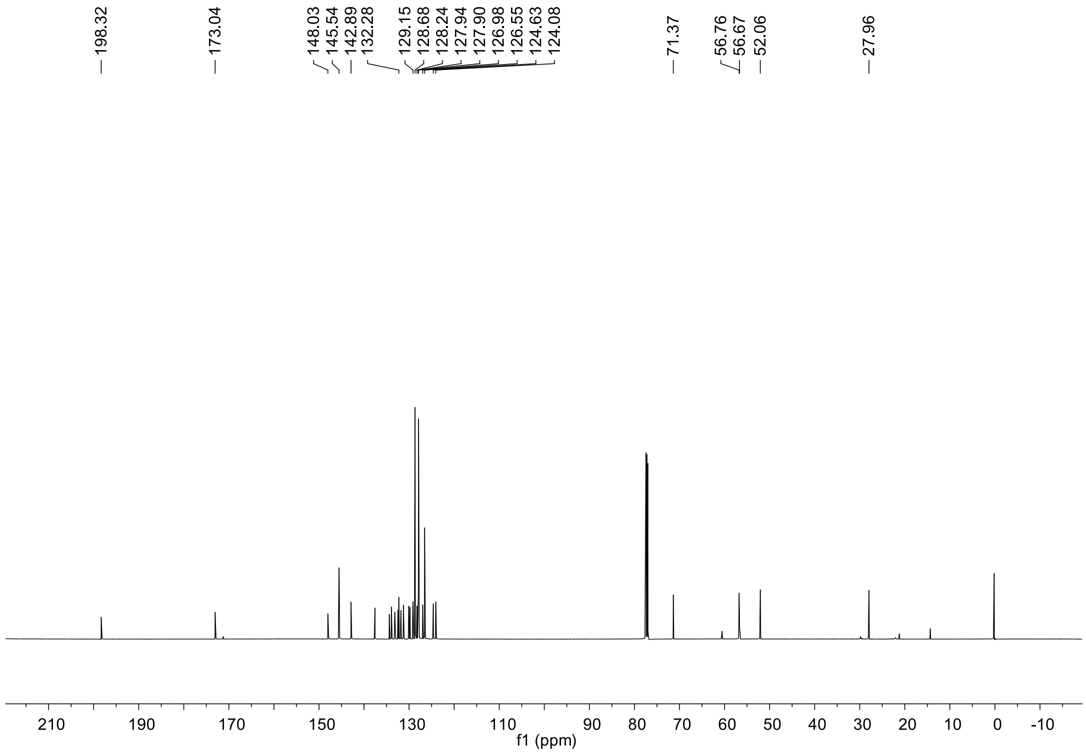

Figure S17. $^{13}$C NMR of compound **20**.

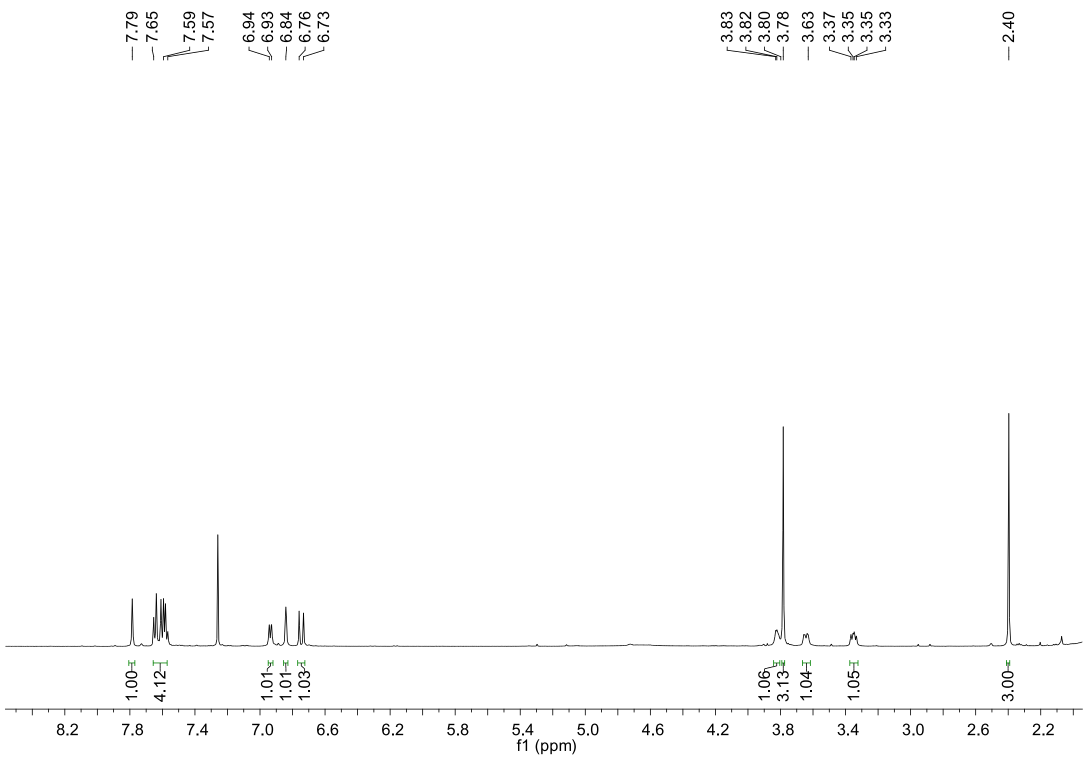

Figure S18. $^{1}$H NMR of compound **22**.

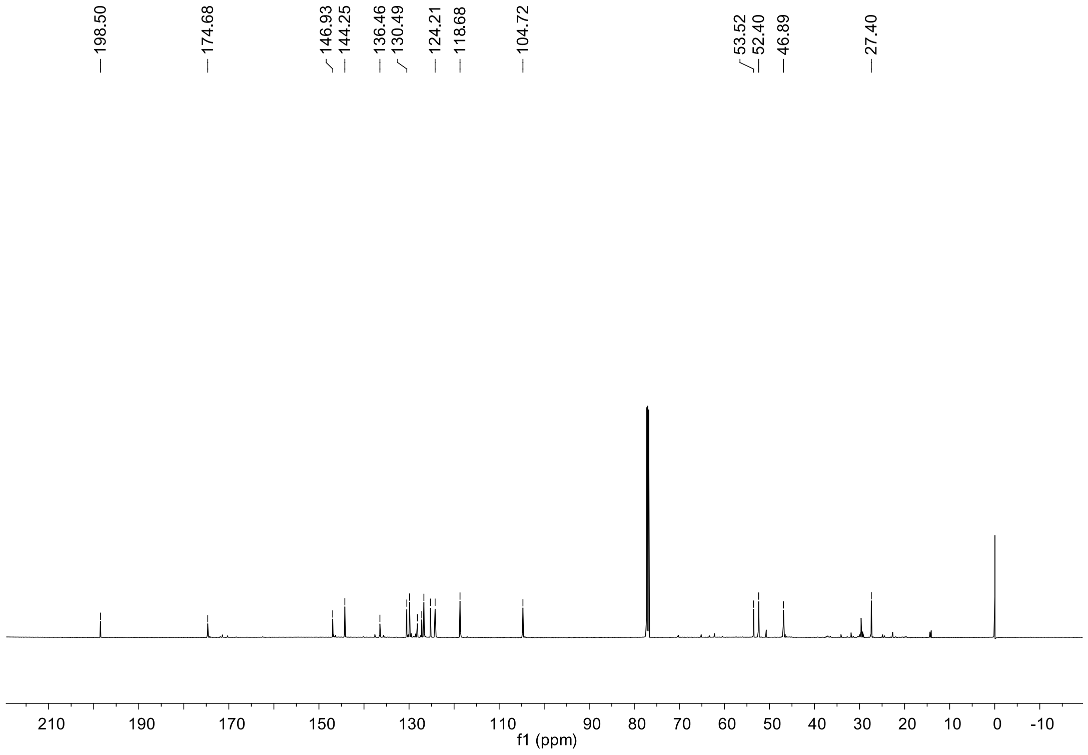

Figure S19. $^{13}$C NMR of compound **22**.

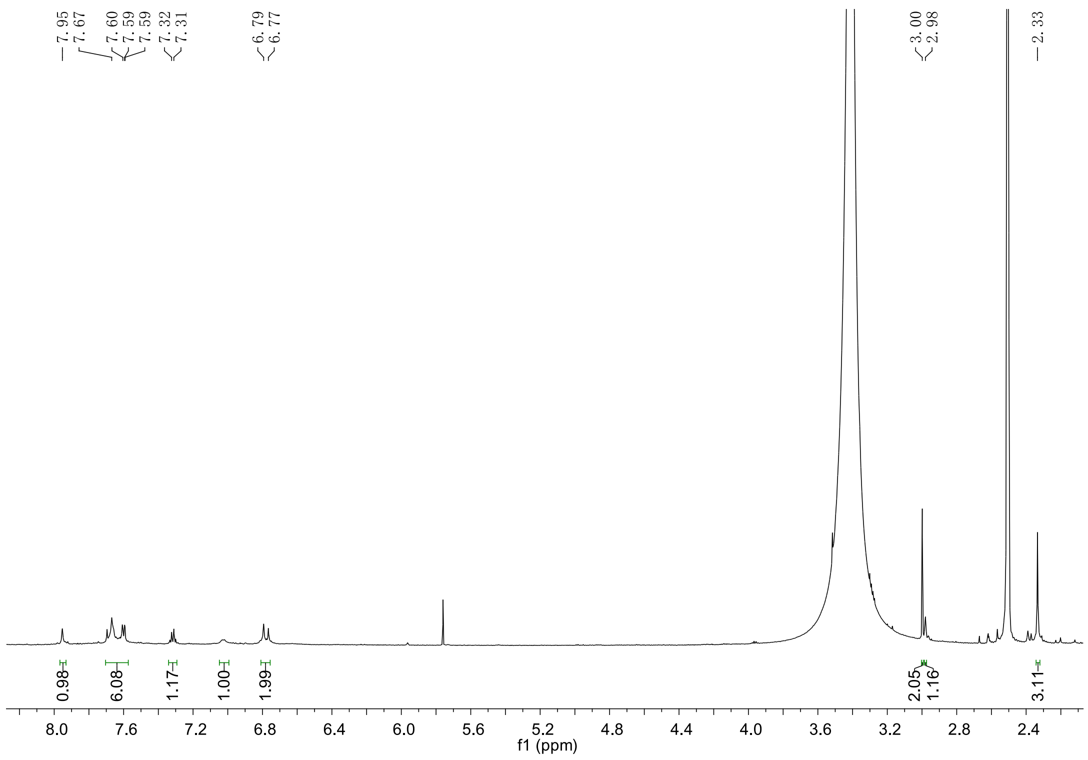

Figure S20. $^1$H NMR of compound **23**.

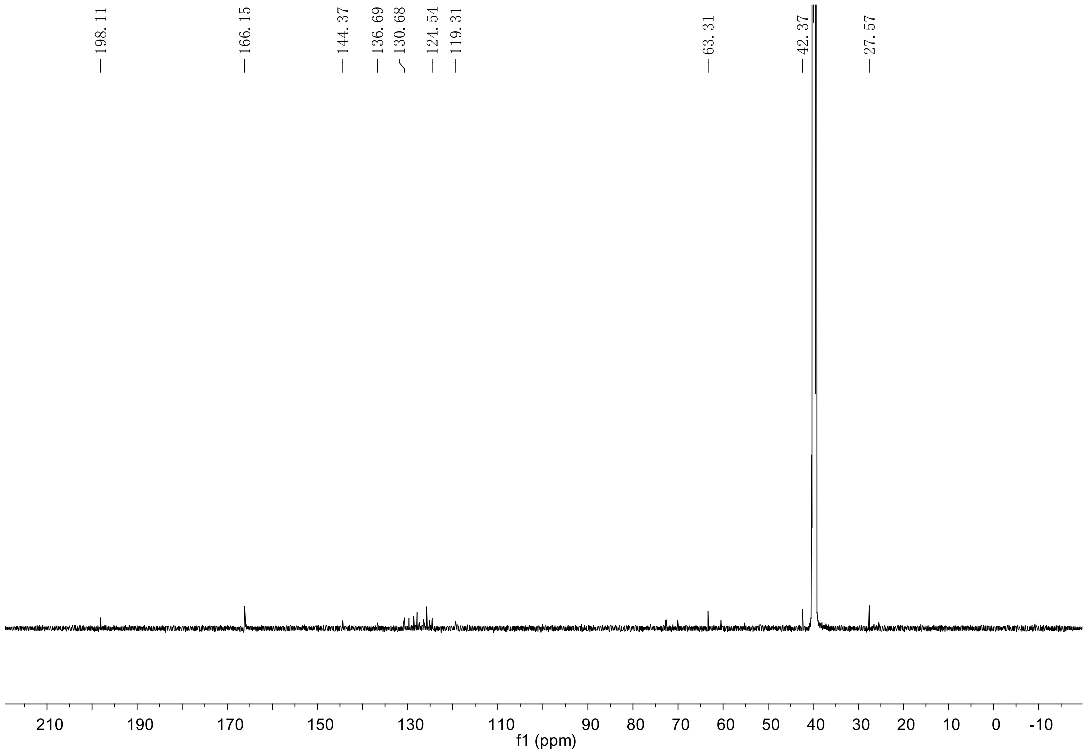

Figure S21. $^{13}$C NMR of compound **23**.

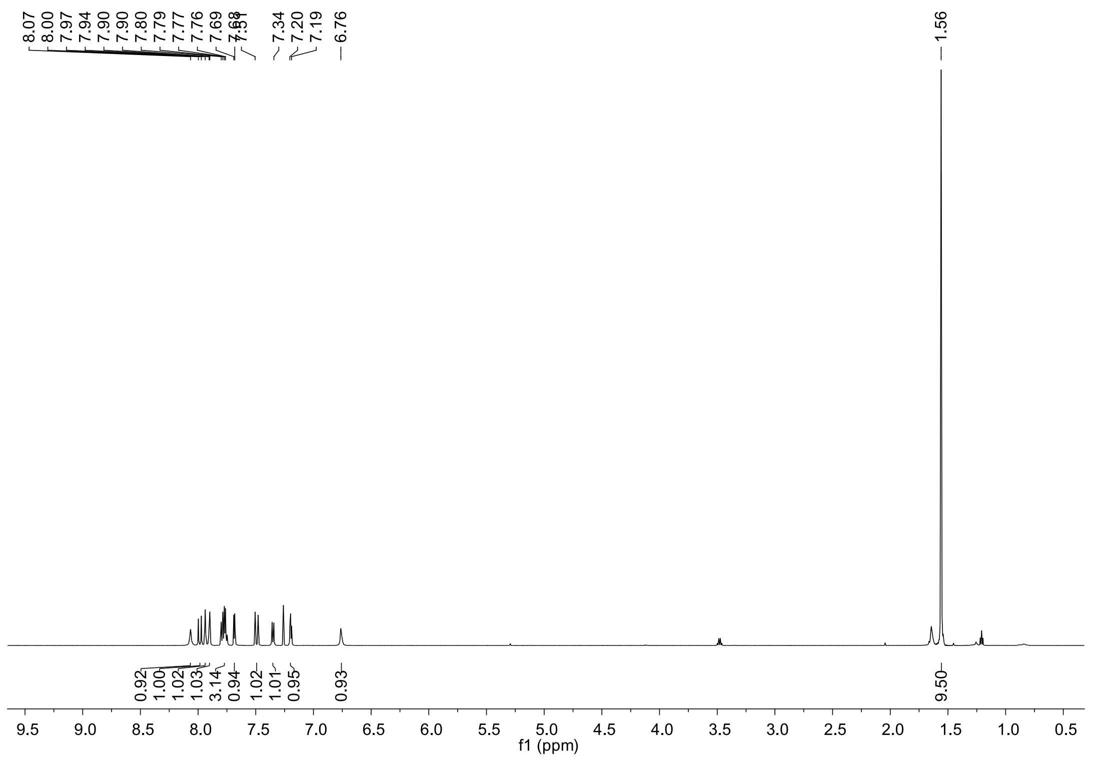

Figure S22. $^{1}$H NMR of compound **24**.

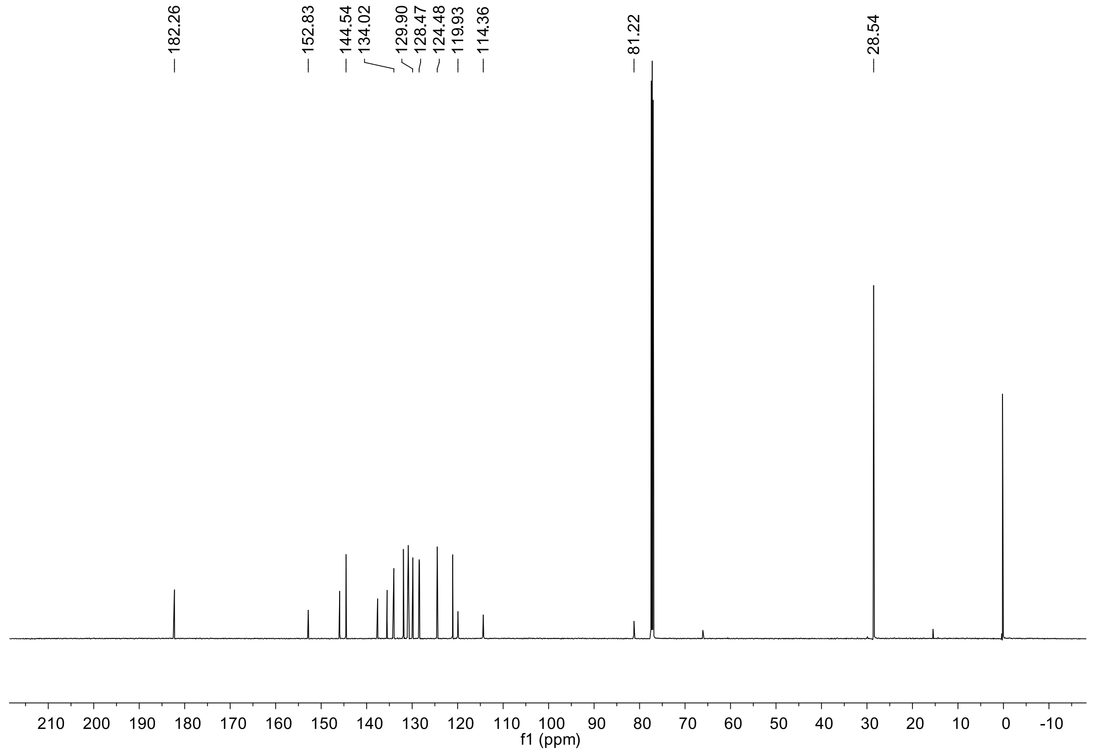

Figure S23. $^{13}$C NMR of compound **24**.

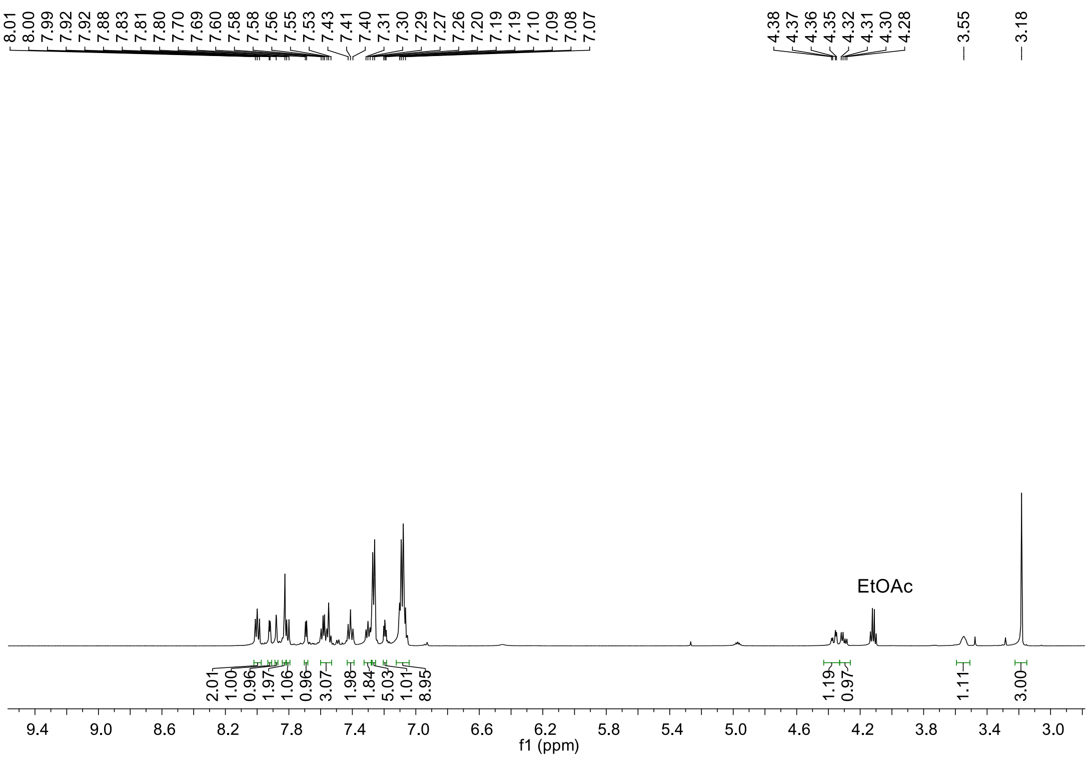

Figure S24. $^{1}$H NMR of compound **27**.

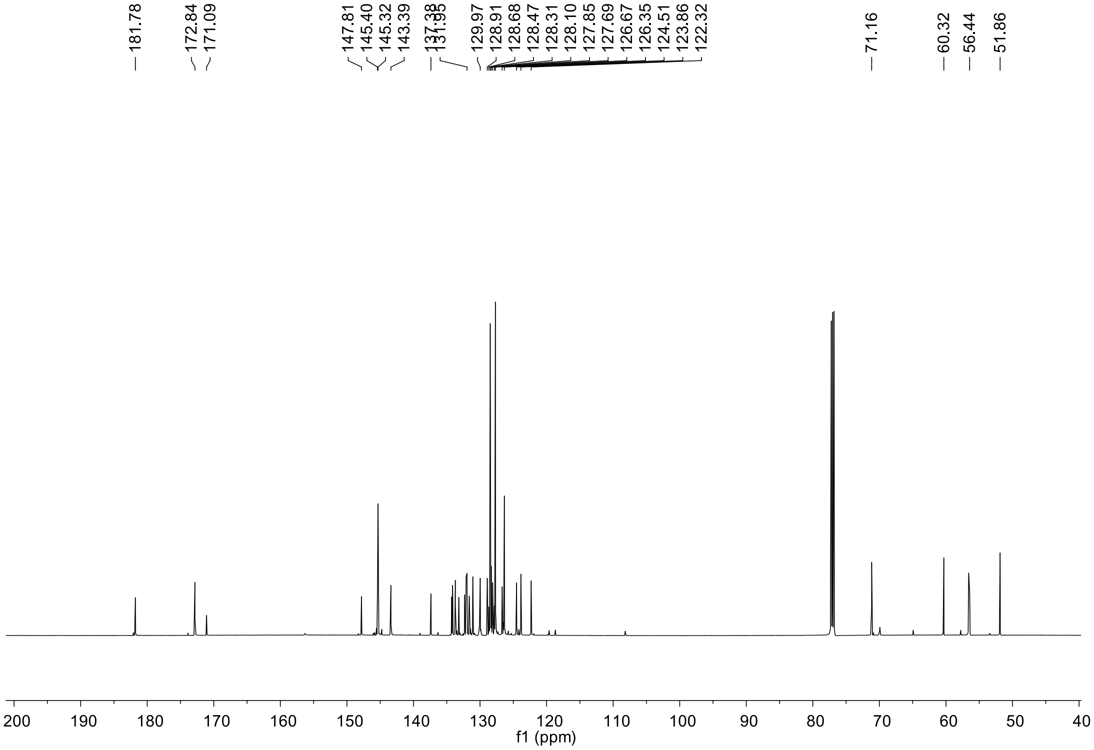

Figure S25. $^{13}$C NMR of compound **27**.

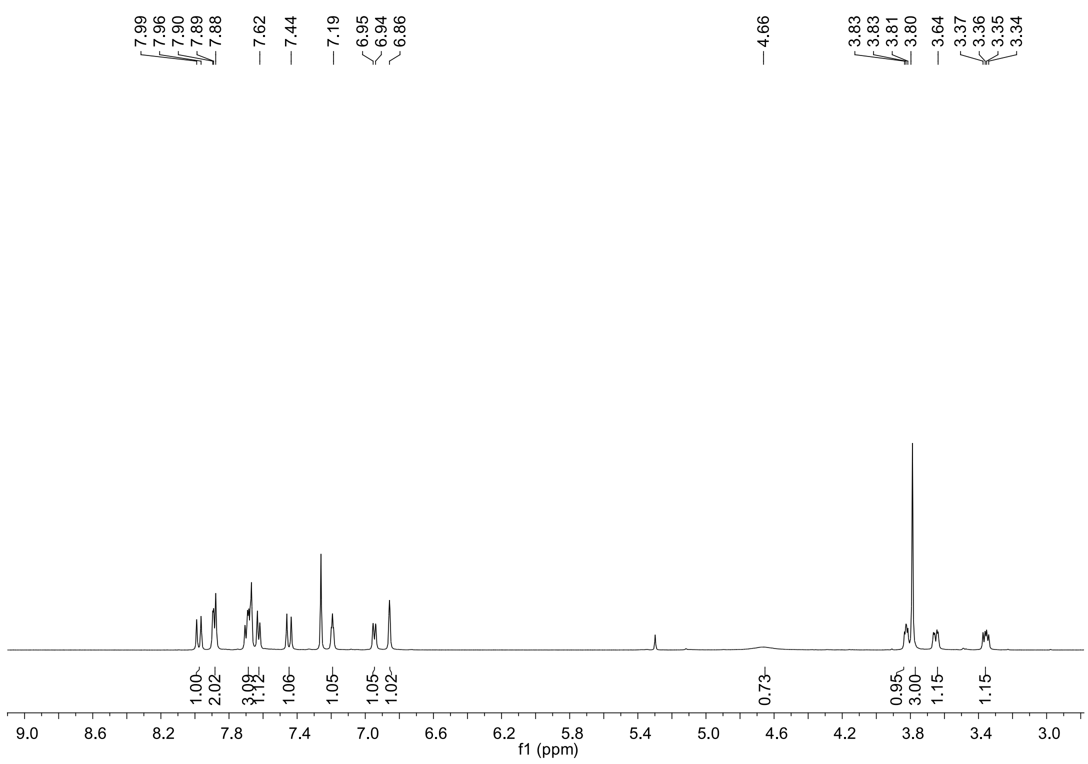

Figure S26. [1]H NMR of compound **29**.

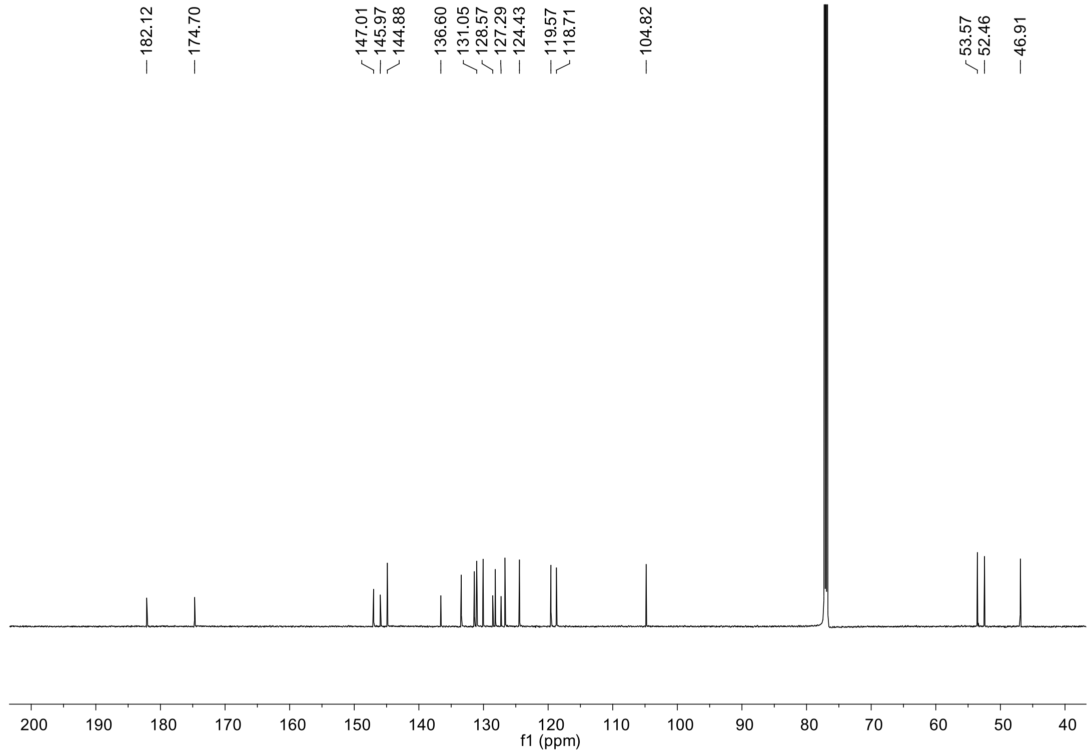

Figure S27. [13]C NMR of compound **29**.

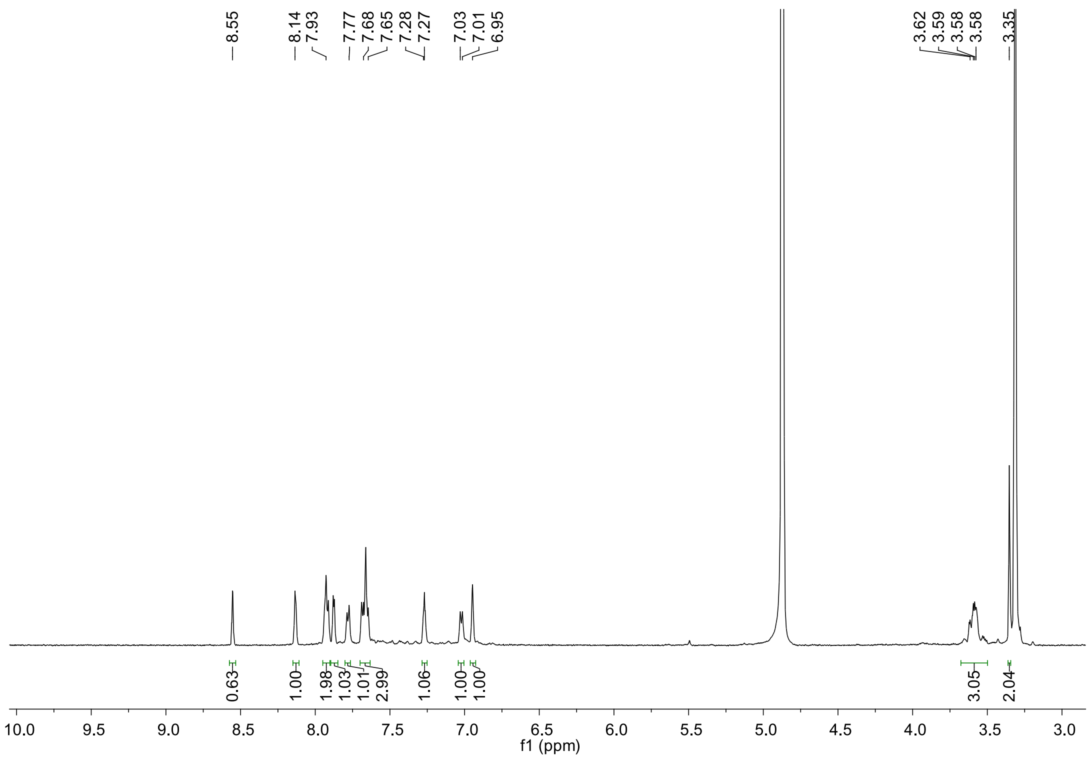

Figure S28. ¹H NMR of compound **30**.

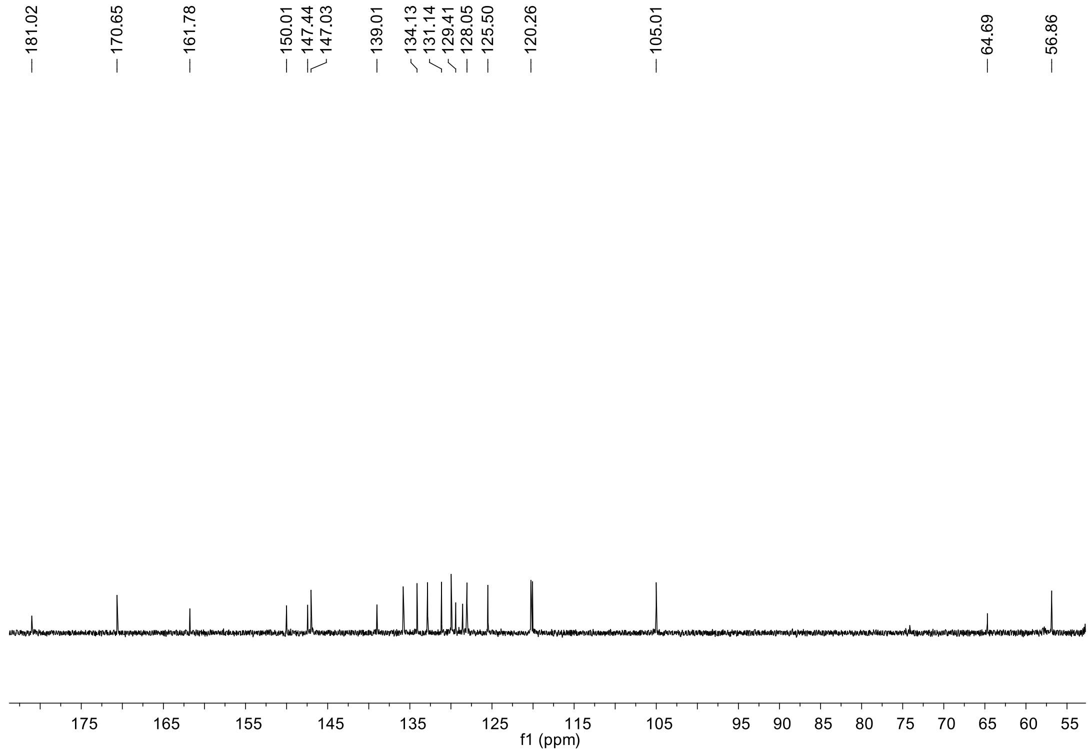

Figure S29. ¹³C NMR of compound **30**.
